# Dominant-negative *TP53* mutations potentiated by the HSF1-regulated proteostasis network

**DOI:** 10.1101/2024.11.01.621414

**Authors:** Stephanie Halim, Rebecca M. Sebastian, Kristi E. Liivak, Jessica E. Patrick, Tiffani Hui, David R. Amici, Andrew O. Giacomelli, Paulina Rios, Vincent L. Butty, William C. Hahn, Francisco J. Sánchez-Rivera, Marc L. Mendillo, Yu-Shan Lin, Matthew D. Shoulders

**Author notes:** Contributed equally to this work (Stephanie Halim, Rebecca M. Sebastian). To whom correspondence may be addressed: Matthew D. Shoulders, Department of Chemistry Massachusetts Institute of Technology 77 Massachusetts Avenue, 16-573A Cambridge, MA 02139.

## Abstract

Protein mutational landscapes are shaped by how amino acid substitutions affect stability and folding or aggregation kinetics. These properties are modulated by cellular proteostasis networks. Heat shock factor 1 (HSF1) is the master regulator of cytosolic and nuclear proteostasis. Chronic HSF1 activity upregulation is a hallmark of cancer cells, potentially because upregulated proteostasis factors facilitate the acquisition and maintenance of oncogenic mutations. Here, we assess how HSF1 activation influences mutational trajectories by which p53 can escape cytotoxic pressure from nutlin-3, an inhibitor of the p53 regulator MDM2. HSF1 activation broadly increases the fitness of dominant-negative p53 substitutions, particularly non-conservative, biophysically unfavorable amino acid changes within buried regions of the p53 DNA-binding domain. These findings demonstrate that HSF1 activation reshapes the oncogenic mutational landscape by preferentially supporting the emergence and persistence of bio-physically disruptive, cancer-associated p53 substitutions, linking proteostasis network activity directly to oncogenic evolution.

## INTRODUCTION

Cancers typically arise by multi-step acquisition of genetic alterations that dysregulate cell growth and survival, leading to malignant phenotypes^1–3^. In addition to mutated genes, cancers also co-opt non-mutated genes within stress pathways to aid cell growth, in a process known as non-oncogene addiction^4, 5^. Pathways associated with maintenance of proteostasis are often upregulated in cancer, likely owing to challenges arising from dysregulated protein synthesis, nutrient starvation, subunit imbalance within protein complexes, and perhaps, expression of oncoproteins with destabilizing amino acid substitutions^6–11^. Most commonly, the heat shock response (HSR), which is controlled by the master transcription factor HSF1, is frequently upregulated in cancer^12, 13^. HSF1 can also modulate additional cell remodeling programs in tumors^14–16^. High basal expression of heat shock proteins regulated by HSF1 is widely observed in malignant cells^13^, as are overexpression and constitutive activity of HSF1 itself^13–15, 17^. HSF1 supports the emergence of tumors in mice following exposure to mutagens, and high levels of HSF1 expression are associated with increased mortality rates in breast cancer^16, 17^. Additionally, HSF1 and the HSF1-regulated chaperone heat shock protein 90 (HSP90) can facilitate the development of resistance to chemotherapeutic agents, although molecular-level mechanisms of this phenomenon are unclear^18, 19^. Accordingly, much attention has been drawn not just to HSP90^20, 21^ and other chaperones but also to HSF1 as a chemotherapeutic target, motivating the development of small molecules targeting HSR components^22, 23^. Although chaperone inhibition has long been implicated as a cancer therapeutic, there are challenges in implementing them as drugs. This challenge suggests that a better understanding between the interplay of chaperones and cancer-associated proteins is needed to successfully translate chaperone modulation into a clinical setting^24^.

One intriguing hypothesis is that an enhanced proteostasis environment amplifies the accessibility of novel mutations in cancer-associated proteins via its influence on the folding and degradation of the resulting protein variants. Successful emergence of gain-of-function mutations in all proteins is constrained by the evolving protein’s biophysical properties, as most functionally important amino acid substitutions are biochemically non-conservative, and therefore, often biophysically deleterious^25, 26^. A growing body of literature has investigated the impact of proteostasis network components on protein evolution^25, 27–34^. Among other advances, such studies have shown that proteostasis network remodeling mediated by stress-responsive transcription factors can have major impacts on the mutational space accessible to viral pathogens that parasitize host chaperones^30–33^, and that these effects are often mediated directly by the influence of proteostasis network composition on client protein folding and stability^25, 28, 32, 35^.

Building on these studies, one compelling possibility is that HSF1 overexpression supports malignancy by creating a permissive protein folding environment that directly facilitates the emergence of proliferation-promoting oncogenic mutations. However, the role for HSF1 in defining accessible cancer protein mutational space has never been experimentally explored.

The tumor suppressor p53 is a key transcription factor and regulator of the response to DNA damage and other oncogenic stimuli. When activated, p53 induces cell-cycle arrest, senescence, or apoptosis^36^. *TP53*, which encodes the p53 protein, is the most frequently mutated gene in cancer^37^. The majority of these mutations are missense mutations within p53’s DNA-binding domain (DBD)^38^. While some *TP53* missense mutations confer a loss-of-function effect, mutations in *TP53* can also function via a dominant-negative effect. In such cases, the resulting p53 variant inhibits the function of residual wild-type p53, possibly through either heterotetramerization via a C-terminal oligomerization domain or by inducing co-aggregation and consequent loss of function^39–41^.

p53 interacts extensively with cellular chaperones, including the HSF1-regulated chaperones HSP90 and HSP70, helping to regulate p53’s stability and activity^42–47^. Notably, while wild-type p53 engages transiently with chaperones, particularly during folding and maturation, several p53 variants display more stable interactions with HSP70 and HSP90, contributing to increased variant p53 levels and variant stabilization^43, 48–51^. Previous work showing that inhibition of HSP90 selectively impacted variants of p53^52, 53^. Mutant p53 aggregation has also been shown to upregulate chaperone levels^54^. Considering the large number of destabilizing p53 substitutions and the extensive interplay between p53 and HSF1-regulated chaperones, p53 represents a compelling model system to explore the impacts of HSF1 activation on the evolution of the cancer-associated proteome.

Here, we apply deep mutational scanning (DMS) of p53 with chemical genetic regulation of HSF1 to examine the impacts of proteostasis modulation on the fitness of dominant-negative p53 variants. We observed that constitutive activation of HSF1 increases the fitness of diverse dominant-negative p53 variants, including several substitutions within hot-spot sites associated with cancer. The impact of HSF1 activation on p53’s mutational spectrum was most evident in destabilizing substitutions of nonpolar to polar amino acids within buried regions of the DBD. These results indicate that HSF1 can directly potentiate oncogene evolution by supporting otherwise biophysically problematic amino acid sequences. Further, these results implicate proteostasis network inhibition as a potential therapeutic strategy to prevent acquisition of resistance mutations during chemotherapy.

## RESULTS

### *TP53* mutational library integrated with chemical genetic regulation of the HSR

To assess the impact of HSF1 on the mutational landscape of dominant-negative p53, we established a system where HSF1 activity could be robustly regulated in an appropriate cell line. We chose to use A549 cells, an epithelial tumor model derived from a human alveolar basal cell adenocarcinoma^55^. A549 cells exclusively express wild-type p53, critically enabling our experimental workflow^56^. Pharmacologic activation of HSF1 is traditionally achieved via treatment with chaperone inhibitors or toxins like arsenite^57^. Such approaches are not useful here, as they activate HSF1 indirectly by causing massive cellular protein misfolding, and ultimately drive apoptosis via proteostatic overload. Instead, we employed regulated expression of a constitutively active HSF1 variant (cHSF1). We constructed a stable, single-colony A549 cell line where we placed cHSF1 expression under the control of a doxycycline (dox)-responsive promoter^58–61^. In these cells (A549^cHSF1^ cells), treatment with dox activates expression of cHSF1, upregulating HSR-controlled gene expression independent of protein misfolding stress.

To test the activation of HSF1, we treated A549^cHSF1^ cells with dox or vehicle for 24 h then evaluated transcript levels of established HSF1 target genes using qPCR. In our optimized cell line, we observed modest upregulation of *DNAJB1* and *HSPA1A* transcripts during HSF1 activation as compared to vehicle treatment, indicating that our cHSF1 construct was functionally modulating the HSR (**Figure 1A**). Critically, we obtained cells where HSF1 induction upregulated HSR genes to levels that are still well within the physiologically accessible regime, to avoid off-target induction of genes not normally targeted by HSF1^60, 62^. As evidence, treatment with STA-9090^63^, an HSP90 inhibitor and robust activator of endogenous HSF1, had considerably stronger effects than the dox treatment (**Figure 1A**). Moreover, a resazurin metabolic activity assay^64^ indicated that cell growth and viability were not altered by dox-mediated HSF1 induction (**Figure S1A**).

**Figure 1:**
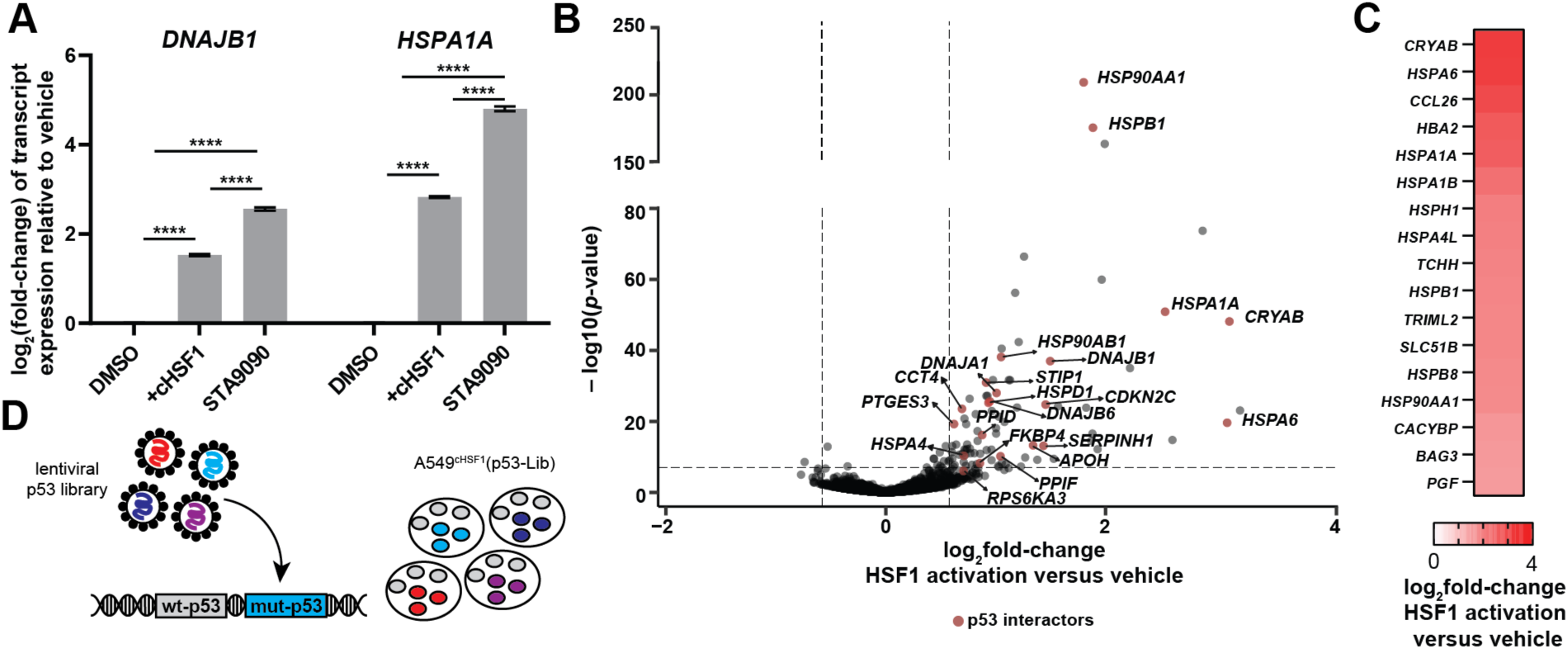
Characterization of chemical genetic HSF1 regulation and construction of the A549^cHSF1^(p53-Lib) cell line. (**A**) qPCR results showing transcript-level consequences of dox-mediated HSF1 activation for the HSF1 target genes *DNAJB1* and *HSPA1A* in A549^cHSF1^ cells. Statistical significance was calculated using two-tailed students *t*-test where **** represents a *p*-value of <0.0001. HSP90 inhibitor STA-9090 was used as a positive control for HSR activation, ensuring activation within the regime accessible to endogenous HSF1 activity. (**B**) Volcano plot for RNA-Seq analysis of changes in gene transcription following HSF1 activation in A549^cHSF1^ cells as compared to vehicle treatment. Proteins within the Agile Protein Interaction DataAnalyzer (APID) that have been identified as interactors with p53 are labeled and shown in red. (**C**) Heat map depicting log_2_ fold-change of transcript expression of selected p53 interacting genes highlighted in (**B**). (**D**) Creation of A549^cHSF1^(p53-Lib) cells: A549 cells expressing endogenous wild-type p53 were transduced at a low multiplicity of infection with a lentiviral population encoding all possible single amino acid substitutions within *TP53.* See also **Table S1** and **S2,** and **Figure S1**.

We aimed to comprehensively assess how HSF1 activation remodeled the proteostasis network. We treated A549^cHSF1^ cells with dox for 24 h and quantified differentially transcribed genes using RNA-seq. 159 transcripts were significantly and differentially expressed with >1.5 log_2_fold-change upon HSF1 activation as compared to the vehicle-treated control, highlighting that HSF1 activation did not massively perturb the global transcriptome (**Table S1**). Known components of the HSR, including *HSP90AA1* and *HSPA1A*, were highly enriched among the upregulated transcripts (**Figure 1B**). Gene set enrichment analysis using the MSigDB c5 collection^65^ further confirmed that genes related to the HSR gene enrichment following activation of HSF1 (**Table S1**). Notably, known p53-induced genes, such as *BAX* and *CDKN1A*, were not substantially impacted by HSF1 activation (**Table S1**).

We asked how HSF1-regulated chaperones might directly alter p53 proteostasis. We identified the subset of transcripts encoding HSF1-regulated chaperones known to interact with p53. Of the 159 transcripts significantly upregulated by HSF1 activation, 22 (13.8%) encode proteins classified by the APID (Agile Protein Interaction DataAnalyzer) database as interacting with p53 (**Table S2**)^66^. Included within these p53-interacting proteins, 17 are either chaperones or co-chaperones, including the well-validated and HSF1-regulated p53 chaperones HSP90 and HSP70 (**Figure 1C**).

We transduced these A549^cHSF1^ cells with a high-quality, lentiviral-based *TP53* mutational library containing all possible single amino acid substitutions across the p53 protein^39, 67^. We transduced at a low multiplicity of infection (MOI = 0.25) to ensure that a single p53 variant was expressed in each cell, *<u>alongside</u>* endogenous wild-type p53 (**Figure 1D**). To evaluate the diversity of the resulting A549^cHSF1^ p53 variant library, (A549^cHSF1^(p53-Lib) cells), we amplified the library-encoded *TP*53 gene using PCR before deep-sequencing. Out of 7467 potential single amino acid substitutions, 7465 (99.9%) of the possible substitutions were observed with a read depth >10 counts per amino acid substitution (**Figure S1B** and **Table S3**).

### HSF1 activation modulates the p53 mutational landscape during MDM2 inhibition

We applied DMS to test whether HSF1 activity alters tolerance for dominant-negative p53 mutations. Our approach was to perform selections in the HSF1-enhanced or basal proteostasis environment in the presence of nutlin-3^67^. Nutlin-3 inhibits the interaction between p53 and mouse double minute 2 homolog (MDM2), an E3 ubiquitin ligase and negative regulator of p53^68^. In unstressed cells, MDM2 binds p53 triggering ubiquitination and degradation. Binding of nutlin-3 to MDM2 releases p53 from the MDM2 complex, allowing p53 to accumulate and induce transcriptional programs driving apoptosis and cell cycle arrest. Cells expressing a dominant-negative p53 variant alongside endogenous wild-type p53 are strongly positively selected in the presence of nutlin-3, because dominant-negative p53 attenuates the wild-type p53-mediated activation of cell cycle arrest and thereby allows cell proliferation (**Figure 2A**)^67^.

**Figure 2:**
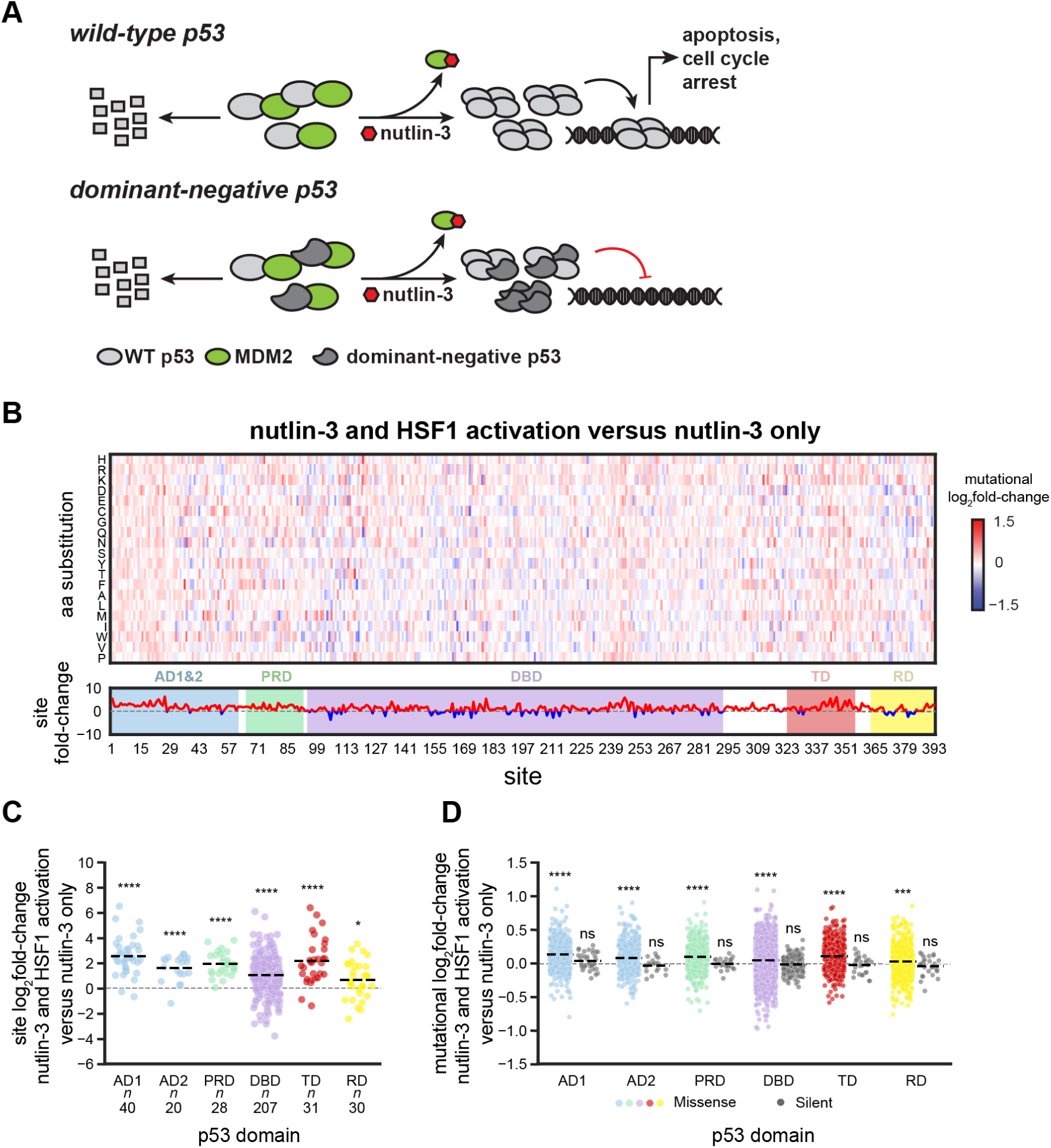
HSF1 activation enhances mutational fitness of dominant-negative p53 variants. (**A**) Selection of dominant-negative variants using nutlin-3. (**B**) Heat map of p53 mutational frequency log_2_ fold-change averaged over three biological replicates for each amino acid substitution for nutlin-3-mediated dominant-negative p53 selection in an HSF1-activated environment as compared to nutlin-3-mediated dominant-negative p53 selection in a basal proteostasis environment (*top*). Sum of the mutational log_2_ fold-change at each site. Color indicates net positive (*red*) or negative (*blue*) site log_2_ fold-change (*bottom*). (**C**) Total site log_2_ fold-change (all substitutions for a given site) for amino acid substitutions in the selection conditions in (**B**) subdivided across each p53 domain. (**D**) Log_2_ fold-change for each individual DNA-level mutation in the *TP53* gene subdivided by domain. Missense mutations (*colored*), synonymous mutations (*grey*). Significance was calculated using a Wilcoxon signed-rank test, with *, ***, and **** representing adjusted two-tailed *p*-values of <0.05, <0.001, and <0.0001, respectively, and ns indicating non-significant. AD1, activation domain 1; AD2, activation domain 2; PRD, proline-rich domain; DBD, DNA-binding domain; TD, tetramerization domain; RD, regulatory domain. See also **Table S3** and **Figures S2–4**.

We treated A549^cHSF1^(p53-Lib) cells with nutlin-3 (or vehicle) in the context of a basal or an HSF1-activated proteostasis environment, allowing the selection to proceed over a 12-day period^67^. After selection, we sequenced the library-encoded *TP53* amplicons, as previously described^67^. We calculated the resulting changes in p53 variant frequency as the log_2_ fold-change in normalized read counts of amino acid substitutions between our various selections versus control conditions (**Table S3**).

We expected that nutlin-3 would be the dominant force affecting the fitness of p53 variants, as cells that fail to express a dominant-negative p53 variant cannot survive nutlin-3 selection. Indeed, we observed a very strong and positive enrichment of missense mutations within the DBD, located from residues 100–300, upon nutlin-3 selection in both the basal (**Figure S2A**) and the HSF1-activated (**Figure S2B**) proteostasis environments relative to the corresponding vehicle-treated (no nutlin-3) controls. Importantly, the sites enriched during this nutlin-3 selection in both the basal and HSF1-activated environments overlapped with previous selections for dominant-negative p53 using saturation mutagene-sis^39, 67^. Moreover, the fitness of p53 variants identified as somatic mutations within the *TP53* data-base^69, 70^ was higher as compared to all missense mutations observed in the DMS experiment, confirming that nutlin-3 specifically selects for cancer-associated, dominant-negative *TP53* mutations (**Figures S2C** and **S2D**). We also observed strong correlations for site log_2_ fold-change values between biological replicates of nutlin-3 treatment versus vehicle under both basal and HSF1 activated proteostasis environments (**Figures S3A** and **S3B**). This observation further indicates that nutlin-3 imposes a very strong selection pressure.

To isolate the impact of HSF1 activation on dominant-negative p53 mutational tolerance, we evaluated whether and how HSF1 activation affected p53 variant fitness specifically during MDM2 inhibition. We observed an increase in p53 variant fitness across much of the *TP53* gene during HSF1 activation (**Figure 2B**). The correlation between individual replicates (**Figure S4A**), was positive and highly significant, indicating the results were reproducible. While reasonable for a DMS experiment^31–33, 71^, the correlation was not as strong as the correlation observed for nutlin-3 treatment versus control (**Figures S3A** and **S3B**), consistent with the stronger selection pressure imposed by nutlin-3 alone.

The observation that chronic HSF1 activation, which is commonly observed across diverse cancers^12–17^, broadly increased p53 mutational tolerance (**Figure 2B**) motivated us to further analyze the underlying effect. We subdivided variants by domain and examined the distribution of the net site log_2_ fold-change and mutational log_2_ fold-change. In all domains, we observed a significant increase in net site fitness (**Figure 2C**). Across these domains we observed an HSF1-dependent increase in mutational fitness for missense mutations only (**Figure 2D**). HSF1 had no significant impact on synonymous mutations (**Figure 2D**), suggesting that the effects of HSF1 are not arising from a general increase in cell fitness. Also noteworthy, HSF1 had minimal effects on p53 variant enrichment in the absence of nutlin-3 treatment, which provides the underlying need for p53 variant selection with MDM2 inhibition (**Figure S4B**). In sum, HSF1 activation specifically enhanced the fitness of non-synonymous dominant-negative *TP53* mutations under MDM2 inhibition, while not impacting synonymous mutations in all domains of p53.

### HSF1 activation supports the accumulation of biophysically destabilizing, non-conservative amino acid substitutions within buried regions of the p53 DNA binding domain

We examined the impacts of HSF1 activation during MDM2 inhibition on variant fitness within the DBD (**Figures 3** and **S5**), because the majority of known oncogenic *TP53* mutations are localized to this region^69^. Moreover, the DBD is the only structurally characterized region of p53. The DBD is composed of a core β-sandwich scaffold supporting a DNA-binding surface comprising two loops (Loop 2 and Loop 3) stabilized by Zn^2+^ coordination, as well as a loop-sheet-helix motif that contains Loop 1^72^.

**Figure 3:**
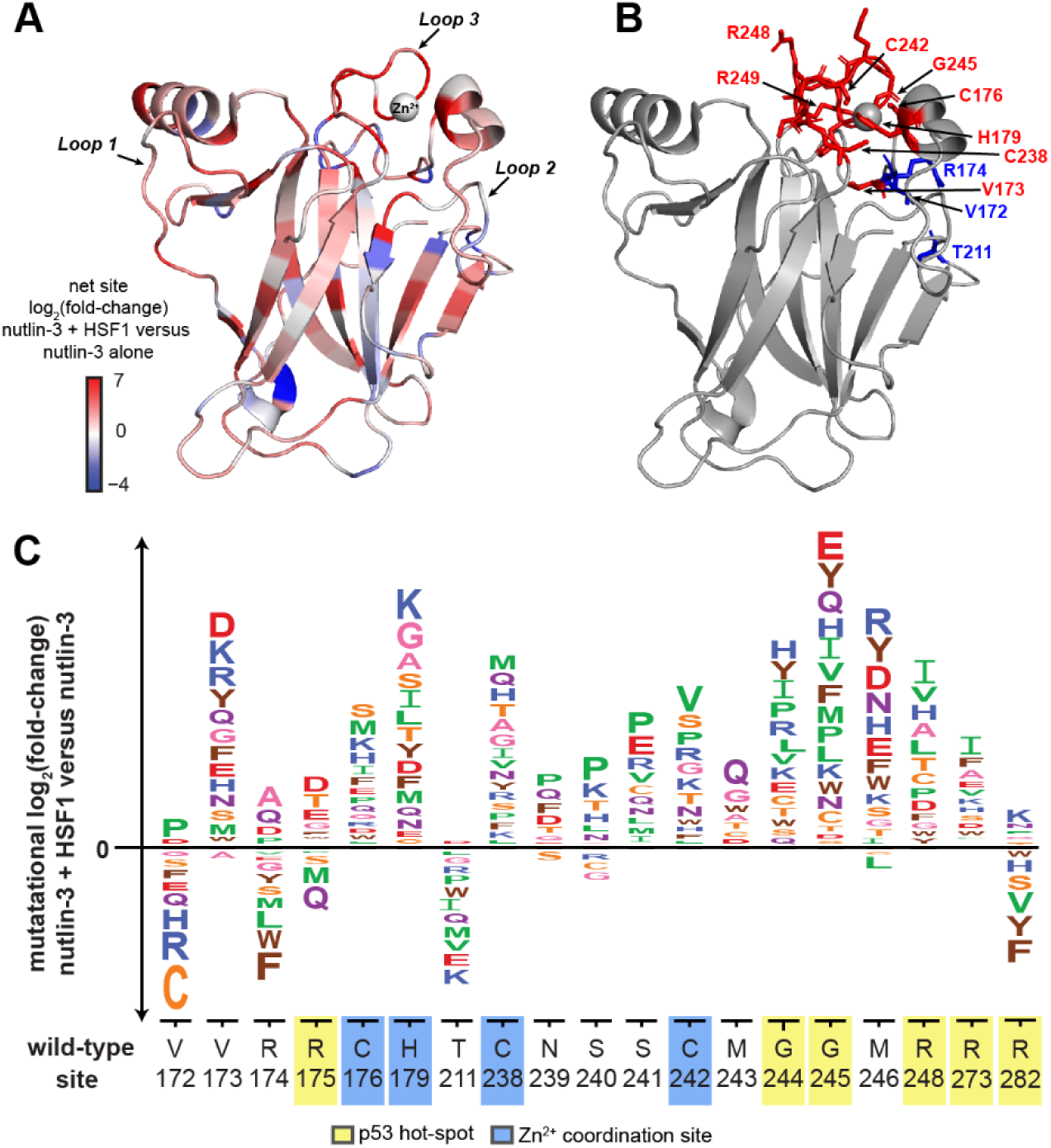
HSF1 activation in the context of nutlin-3-mediated selection alters mutational fitness within the p53 DNA-binding domain. (**A**) Structure of the p53 DNA-binding domain (PDB 2OCJ). Residues are colored according to the net site log_2_ fold-change. (**B**) Consequences of HSF1 activation at selected, individually labeled hot-spot sites. Color indicates net positive (*red*) or negative (*blue*) site log_2_ fold-change. (**C**) Sequence logo plot of mutational log_2_ fold-change for selected sites, full DBD sequence logo plot available in **Figure S5**. Alleles displaying opposing signs between biological replicates were removed. p53 hot-spot sites are highlighted in *yellow* and Zn^2+^ coordination sites are highlighted in *blue*.

The impact of HSF1 was seen most in sites 238–249 within Loop 3, and sites V173 and H179 within Loop 2 (**Figure S5**). Several cancer hot-spots are localized in this region, including sites G245, R248, and R249, all of which displayed a net increase in fitness upon HSF1 activation. These observations indicate that HSF1 can directly support the acquisition of hot-spot p53 mutations associated with malignant transformation.

Relatively fewer p53 sites displayed an overall net negative fitness upon HSF1 activation during MDM2 inhibition. Interestingly, several of the few sites with net negative fitness (V172, R174, and T211; **Figures 3B** and **3C**) are at the surface of a pocket that interacts with the N-terminal tail of the DBD. Interaction of the N-terminal tail of the DBD with residues in this pocket has been shown to both increase p53 thermodynamic stability as well as decrease aggregation propensity^73^. A possible explanation for the decrease in fitness at these sites is that variants within this region increase the propensity for mutant p53 to co-aggregate with wild-type p53, with resultant dominant-negative consequences favorable in nutlin-3 selection. Such destabilization and aggregation may be attenuated in the supportive proteostasis environment created by HSF1 activation, leading to their relative decrease in fitness during simultaneous HSF1 activation and MDM2 inhibition.

Prototypical oncogenic p53 amino acid substitutions are classed into either DNA-contact variants with minimal impact on thermodynamic stability or structural mutations that significantly perturb DBD stability^74^. Mutations within this second class are frequently localized to either the Zn^2+^-binding site, or within the β-sandwich motif at the hydrophobic core of the DBD, prompting us to analyze these sites in more detail. In particular, Zn^2+^ coordination is critical for p53’s DNA binding ability, and loss of Zn^2+^ binding is associated with destabilization and aggregation of p53^74–76^. We examined the fitness of amino acid substitutions within the Zn^2+^ coordination sites (C176 and H179 of Loop 2 and C238 and C242 of Loop 3; **Figures 3B** and **3C**). As with other cancer-associated mutations, we observed that the net site fitness for all four coordinating residues increased following HSF1 activation (**Figure 3C**).

Substitutions within the β-sandwich motif and at the hydrophobic core of the DBD could be particularly biophysically disruptive. We tested whether there was a correlation between relative solvent accessibility (RSA; **Table S4**) and net site fitness within the DBD because of nutlin-3-mediated p53 selection regardless of the proteostasis environment. We observed a negative correlation between net site fitness and RSA during nutlin-3 selection versus vehicle treatment in either the basal or in the HSF1-activated proteostasis environment (**Figures S6A–D**). This observation coincides with the expectation that substitutions to buried residues within the p53 DNA binding domain are likely to elicit domi-nant-negative behavior and merit further scrutiny.

Accordingly, we asked whether HSF1 activation during MDM2 inhibition preferentially impacted the fitness of p53 amino acid substitutions within buried regions of the DBD. We noted that variants classified as buried (RSA < 0.2) displayed marginally higher net fitness as compared to sites that were exposed (RSA > 0.2; **Figure S6E**). Motivated by this result, we clustered variants into three classes based on their biochemical properties:

a. Conservative substitutions (nonpolar amino acid → nonpolar amino acid or polar → polar amino acid).
b. Non-conservative substitutions (nonpolar →polar amino acid).
c. Non-conservative substitutions (polar → nonpolar amino acid).

We analyzed whether HSF1 activation impacted non-conservative substitutions more strongly than conservative substitutions. In exposed DBD regions, there was no significant effect in any mutation class upon HSF1 activation. However, non-conservative substitutions replacing a nonpolar amino acid with a polar amino acid within buried p53 sites displayed substantially and significantly higher fitness upon HSF1 activation relative to the other two substitution classes (**Figure 4A**). These results suggest that HSF1 activation particularly enables p53 to more robustly access non-conservative, bio-physically disruptive substitutions — especially the introduction of polar residues in hydrophobic, buried domains.

**Figure 4:**
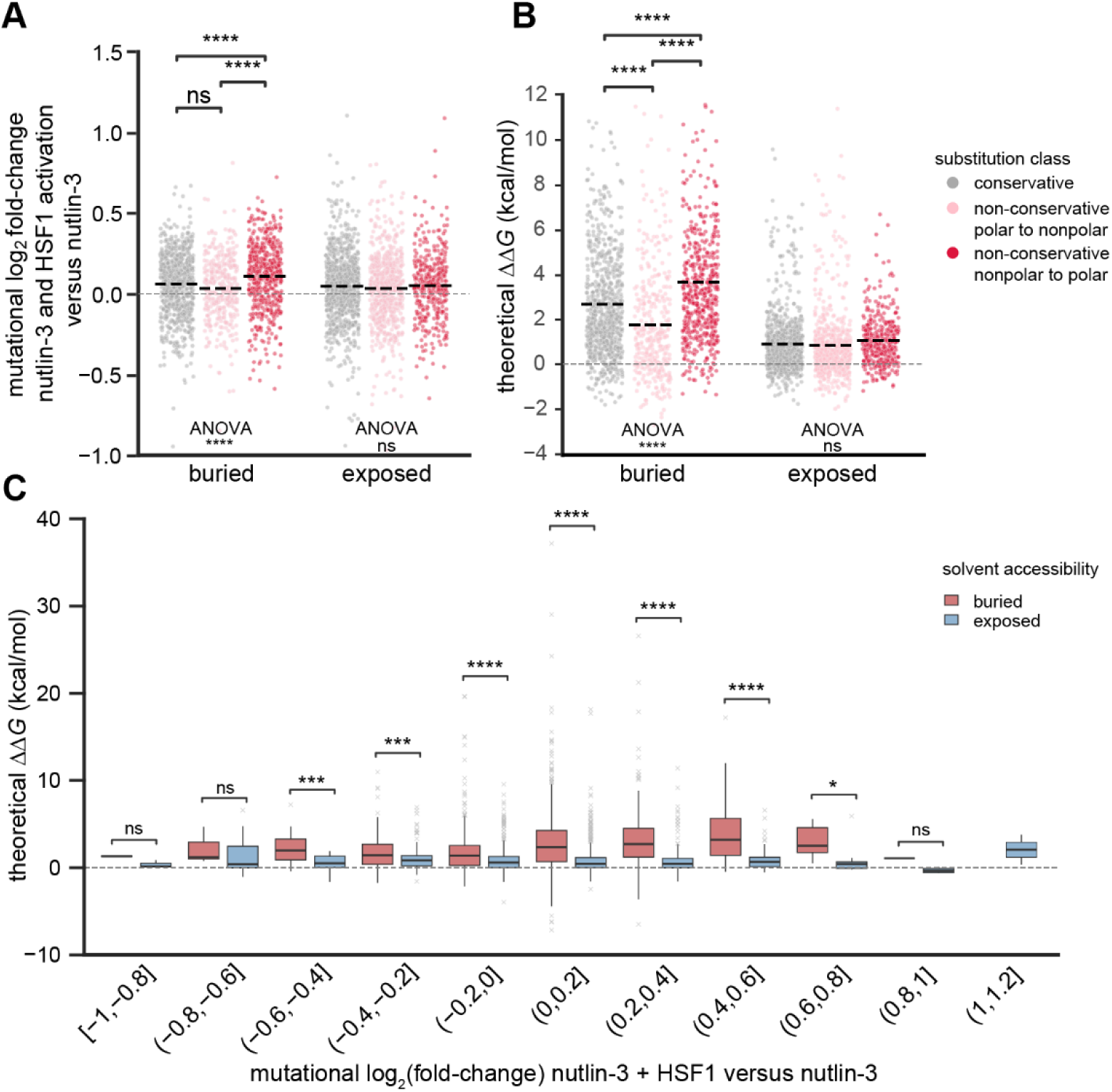
HSF1 most strongly increases fitness of destabilizing substitutions involving biophysi-cally disruptive replacement of nonpolar residues with polar in buried regions of the DBD. (**A**) Mutational log_2_ fold-change in variant fitness for conservative amino acid substitutions, non-conserva-tive nonpolar to polar substitutions, and non-conservative polar to nonpolar substitutions at buried (RSA < 0.2) versus exposed (RSA > 0.2) sites in the p53 DNA-binding domain. (**B**) Theoretical ΔΔ*G* calculated using Rosetta analysis for p53 DNA-binding domain substitutions in buried or exposed sites. For (**A**) and (**B**), statistical significance between solvent accessibility classes or mutation types within a solvent ac-cessibility class was evaluated using ANOVA, while comparisons between select conditions were calcu-lated using Welch’s *t*-test for independent samples with Bonferroni correction. *, ***, and **** represent adjusted two-tailed *p*-values of <0.05, <0.001, and <0.0001, respectively (**C**) Theoretical ΔΔ*G* of buried and exposed variants binned according to mutational log_2_ fold-change in HSF1-activated versus basal proteostasis environments during nutlin-3 selection. Individual box plots represent bins of 0.2 mutational log_2_ fold-change greater than the lower limit and up to and including the upper limit. Outliers are repre-sented by grey crosses. Statistical significance was calculated using a Wilcoxon signed-rank test, with *, ***, and **** representing adjusted two-tailed *p*-values of <0.05, <0.001, and <0.0001, respectively. See also **Figure S6** and **Tables S4** and **S5**

To further explore this phenomenon, we examined 42 missense mutations in the DBD for which thermodynamic stability has been experimentally determined^74, 77–79^. We grouped the variants into those that were stabilizing (ΔΔ*G* < 0.125 kcal/mol) or destabilizing (ΔΔ*G* > 0.125 kcal/mol). We observed an increase in fitness for mutations that were thermodynamically destabilizing as compared to stabilizing mutations (**Figure S6F**).

Since thermodynamic stability measurements have been experimentally performed for only a very limited number of DBD variants and were reported by several different studies, we built on the trend in **Figure S6F** by using Rosetta to estimate the change in thermodynamic stability for all possible amino acid substitutions within the DBD (**Table S5**)^80^. To assess the accuracy of the calculated ΔΔ*G* values, we compared the computed to the experimentally determined ΔΔ*G* values for the aforementioned 42 amino acid substitutions. We observed a strong correlation (**Figure S6G**), validating the computational approach.

As expected, our computations predicted that substitutions from nonpolar to polar amino acids in buried regions have the greatest impact on p53 stability in buried regions of the DBD (**Figure 4B**). The trend in **Figure 4B** mirrored the trend discussed in **Figure 4A**, namely that HSF1 activation impacted non-conservative nonpolar to polar substitutions in buried regions of the DBD of p53 the most. We assessed whether and how HSF1 activation preferentially impacted thermodynamically destabilizing amino acid substitutions in the DBD. To evaluate whether there was a difference in estimated thermodynamic stability depending on the direction and magnitude of their change in fitness with HSF1 activation in a nutlin-3 environment we binned variants by their mutational fold-change HSF1 activation and nutlin-3 versus nutlin-3 alone selection, then plotted the predicted ΔΔ*G*. In buried regions of the DBD, we observed that substitutions with enhanced fitness upon HSF1 activation were more likely to display a higher ΔΔ*G* than variants that displayed decreased fitness (**Figure 4C**). In contrast, we observed little difference in the effect of HSF1 activation on destabilized versus stabilized variants for exposed sites of the DBD (**Figure 4C**). In sum, HSF1 activation specifically supports the emergence of destabilizing p53 substitutions during MDM2 inhibition.

### HSF1-potentiation of destabilizing and oncogenic dominant-negative p53 variants is reproducible in head-to-head competition assays and generalizable across cancer cell lines

With the key conclusion from our large-scale DMS data established, we pursued detailed studies on some specific p53 variants. We studied two p53 variants (V173Y and F113K) in buried regions of the DBD that were predicted to be biophysically destabilizing (ΔΔ*G* > 5 kcal/mol) and representing the nonpolar-to-polar substitutions we found to be most potentiated by HSF1. F113K is also charged in addition to being polar (**Figure 5A**). We also studied a prototypical oncogenic, dominant-negative p53 substitution commonly found in cancer patients (R273H). p53^R273H^ displayed enhanced fitness upon HSF1 activation in our DMS experiments, albeit with a smaller effect size than the F113K or V173Y substitutions.

**Figure 5:**
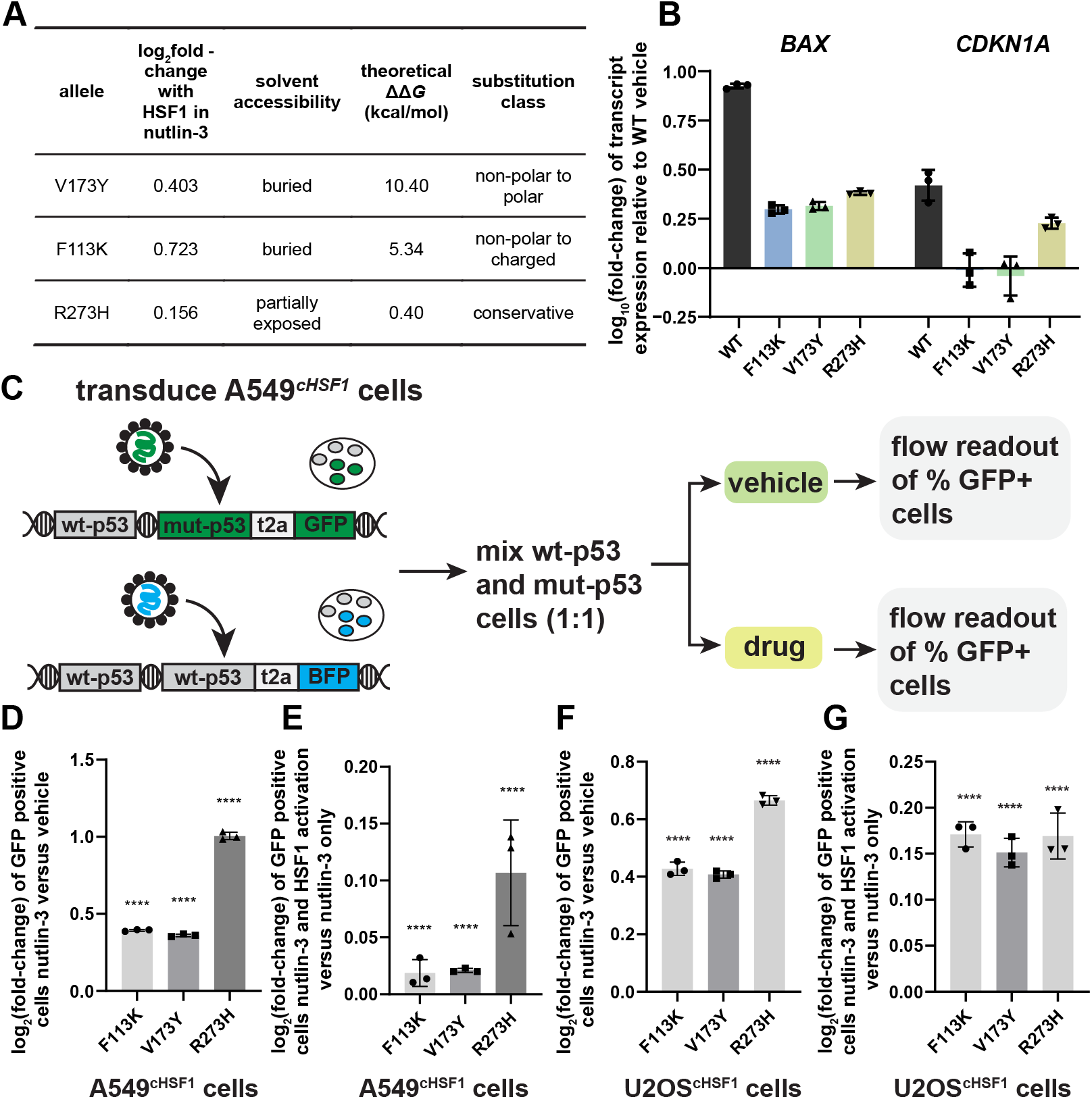
HSF1-potentiated dominant-negative p53 variants identified in the DMS experiment increase in fitness upon HSF1 activation in multiple cancer cell lines. (**A**) Summary table of selected p53 variants. (**B**) qPCR results showing transcript expression of p53 target genes *BAX* and *CDKN1A* of cells expressing wild-type p53 and an HSF1-potentiated dominant-negative p53 variant during HSF1 activation and nutlin-3 treatment**. (C)** Workflow of flow cytometry-based pairwise competition assay. Log_2_ fold-change of GFP frequency as a readout of mutant p53 enrichment under nutlin-3 selection versus vehicle (**D**) and nutlin-3 and HSF1 activation versus nutlin-3 only (**E**) in A549^cHSF1^ cells. Log_2_ fold-change of GFP+ cells as a readout of mutant p53 enrichment under nutlin-3 selection versus vehicle (**F**) and nutlin-3 and HSF1 activation versus nutlin-3 only (**G**) in U2OS^cHSF1^ cells. Significance was assessed using Fisher’s exact test for count data, with **** representing an adjusted two-tailed *p*-value <0.0001. See also **Figure S7** and **Table S6.**

We conducted qPCR assays to assess the dominant-negative activity of these p53 variants in an HSF1-activated environment. We observed strong suppression of the transcript levels of p53 target genes, *BAX* and *CDKN1A* (p21), when these variants were co-expressed with wild-type p53 in an HSF1-activated environment alongside MDM2 inhibition via nutlin-3 treatment (**Figure 5B**).

With dominant-negative behavior demonstrated, to test our results in a focused biochemical experiment we performed a pairwise competition experiment for cells expressing only wild-type p53 versus cells expressing both wild-type p53 and each individual p53 variant under the relevant conditions (**Figure 5C**). Briefly, we transduced A549^cHSF1^ cells with lentivirus encoding either wild-type or variant p53 along with a corresponding fluorescent marker: BFP for wild-type p53 or GFP for p53 variants (**Figure 5C**). These markers enabled fluorescent readout of the abundance of each variant in a pre-competition mixture, as well as 72–96 h post-treatment with either nutlin-3 alone or nutlin-3 and dox to induce HSF1. We observed enrichment of the p53 variants relative to wild-type upon nutlin-3 treatment, confirming that the variants are exerting a dominant-negative effect (**Figure 5D**)^67, 68^. Upon HSF1 activation, we observed an additional, significant enrichment of these same variants (**Figure 5E**), validating the DMS results in A549^cHSF1^ cells for individual variants. Notably, the common oncogenic p53 variant R273H showed a larger effect size than the other variants in the head-to-head competition, suggesting that our A549^cHSF1^ DMS results may underestimate the importance of HSF1 activity in potentiating common oncogenic p53 substitutions.

To assess whether the fitness-enhancing consequences of HSF1 activity for dominant-negative p53 substitutions are generalizable beyond A549 cells, we pursued a similar set of competition experiments in U2OS cells, an osteosarcoma cell line^81^. Importantly, U2OS cells, like A549 cells, are a cancer line that genomically encodes wild-type p53^82^. Additionally, the basal transcript HSF1 expression of U2OS cells is higher than in A549 cells (**Figure S7A**). As with our A549 cells, we engineered U2OS^cHSF1^ cells with dox-regulated induction of HSF1 activity (**Figure S7B**). We then introduced the same p53 variants (**Figure 5A**) and repeated the competition experiments shown in **Figure 5C** using the resulting U2OS^cHSF1^ cell lines. Once again, we observed enrichment of the p53 variants relative to wild-type p53 upon nutlin-3 treatment, consistent with these variants exerting a dominant-negative effect (**Figure 5F**) in U2OS cells. This enrichment was strongly potentiated upon HSF1 activation, with even larger effect sizes than in A549^cHSF1^ cells (**Figure 5G**). Thus, the effects of HSF1 activity on p53 variant fitness are generalizable beyond just one cancer model.

### HSF1-potentiation of dominant-negative p53 substitutions is associated with aggregation modulation and likely involves the activities of HSF1-induced chaperones

Amino acid substitutions that induce mild structural defects within the DBD are likely to function through tetramerization with wild-type p53, where the presence of one or more non-functional p53 sub-units reduces the ability of the overall complex to bind to DNA and induce transcription^40, 83^. For such non-functional p53 subunits to retain dominant-negative function, they must be soluble and not aggre-gated in cells. Interestingly along these lines, confocal microscopy analysis of A549^cHSF1^ cells co-expressing wild-type p53 and the V173Y variant revealed visible p53 aggregates in the nucleus under basal conditions (**Figure 6A**). Likely due to the rapid clearance of wild-type 53, we were unable to image wild-type p53 alone. HSF1 activation resulted in reduced aggregation and even total clearance of aggregation in some cells, suggesting that HSF1 may act to prevent aggregation-inducing conformations of destabilized p53 variants in a manner that promotes dominant-negative function via tetramerization (**Figure 6B**).

**Figure 6:**
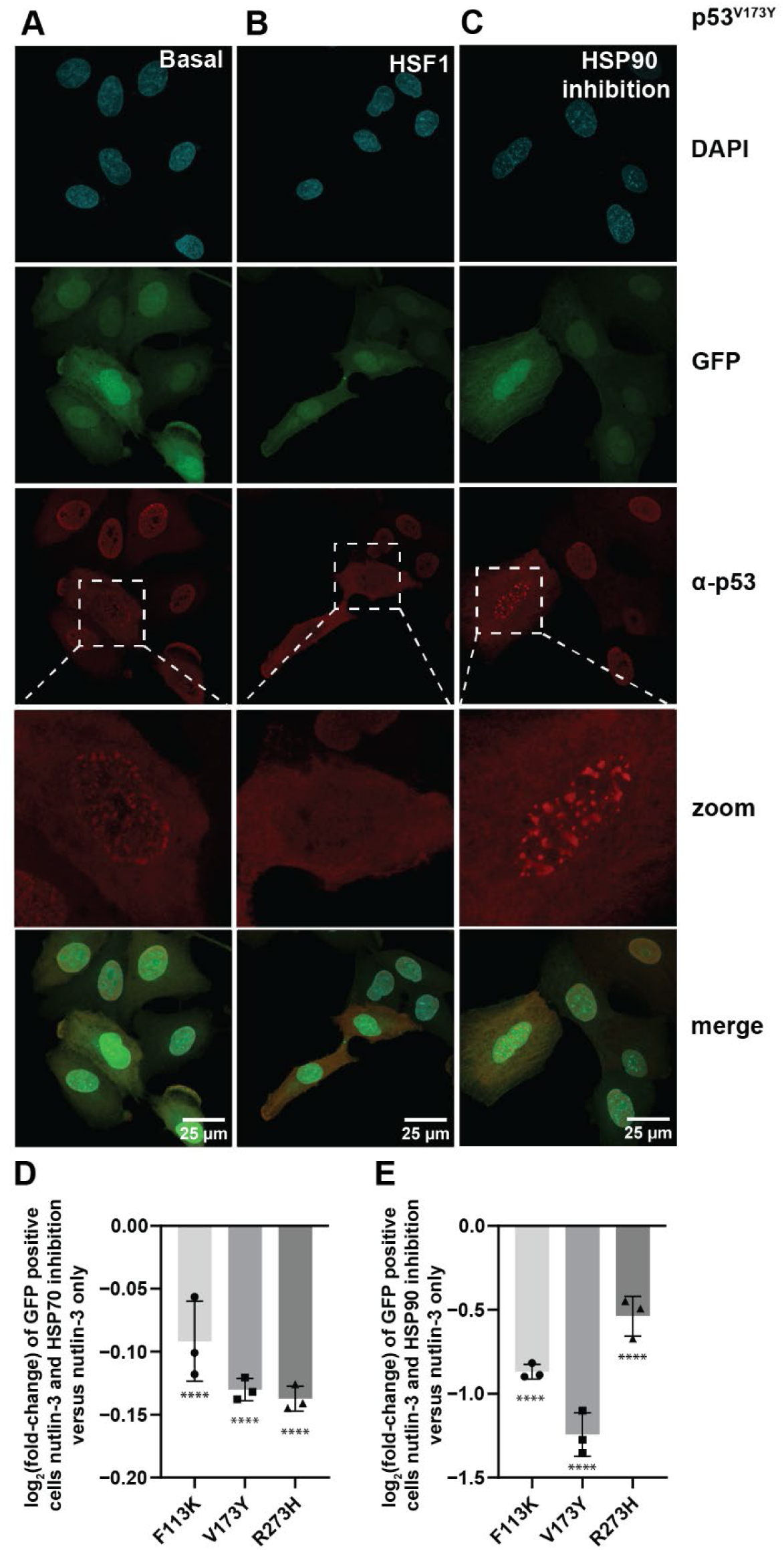
The destabilizing p53 V173Y variant forms aggregates in A549 cells that are cleared by HSF1 activation. Inhibition of chaperones downstream of HSF1 compromises p53 mutant fitness and induces p53 aggregation in cells. A549^cHSF1^ cells stably expressing GFP (to visualize transduction efficiency), wild-type p53, and p53^V173Y^ were treated with either vehicle (**A**), dox to activate HSF1 (**B**), or STA-9090 to inhibit HSP90 (**C**). Selected confocal images of the cells stained for p53 and nuclear stain DAPI reveal that nuclear p53 aggregation under basal conditions was cleared by HSF1 activation while p53 aggregation found in the nucleus was exacerbated with HSP90 inhibition. Log_2_ fold-change of mutant p53 with HSP70 (**D**) and HSP90 (**E**) inhibition in a pairwise competition assay. Significance was calculated using a Fisher’s exact test for count data, with **** representing adjusted two-tailed *p*-values of <0.0001.

We sought to better understand the underlying forces shaping HSF1 potentiation of dominant-negative p53 variants. Specifically, we hypothesized that inhibition of key p53-interacting chaperones induced by HSF1 might reduce the fitness of HSF1-potentiated p53 variants in a nutlin-3 environment. We performed pairwise competition assays with the V173Y, F113K, and R273H p53 variants identified from the DMS screen in the presence of either an HSP90 inhibitor (STA-9090) or an HSP70 inhibitor (VER-155008) in an MDM2-inhibited environment^63, 84^. Based on the potentiation of these variants by HSF1 activation, we hypothesized that they would perform more poorly upon strong chaperone inhibition. We observed a decrease in p53 mutant fitness in the presence of these chaperone inhibitors (**Figures 6D** and **6E**). We asked if the reason for the loss in fitness could be related to excessive mutant p53 protein aggregation. After treating V173Y p53-expressing cells for 24 h with STA-9090 to inhibit HSP90, we again visualized p53 using confocal microscopy. We observed increased V173Y p53 aggregation in some cells upon HSP90 inhibition (**Figure 6C**). Taken together, these observations support the idea that chaperone networks modulate p53 variant fitness by tuning protein stability and aggregation propensity.

### HSF1-potentiated p53 variants are enriched in patients

We questioned whether HSF1 activation impacts p53 mutations observed in cancer patients. We analyzed distributions of the effects of HSF1, represented by the mutational log_2_ fold-change of HSF1 and nutlin-3 versus nutlin-3 alone, for all possible missense mutations from our DMS data to variants observed in cancer patients in the *TP53* database^69, 70^. We compared the mutational fold-change of nutlin-3 and HSF1 versus nutlin-3 alone of all missense mutations in the DMS and missense mutations with and without the dominant-negative loss of function (DNE_LOF) annotation in the *TP53* database (**Figure 7A**). We observed that missense mutations annotated as DNE_LOF were significantly impacted by HSF1 activation in a nutlin-3 environment, while missense mutations without the DNE_LOF annotation were not potentiated by HSF1. This comparison suggests that HSF1-potentiated dominant-negative p53 variants are enriched in the patient population.

**Figure 7:**
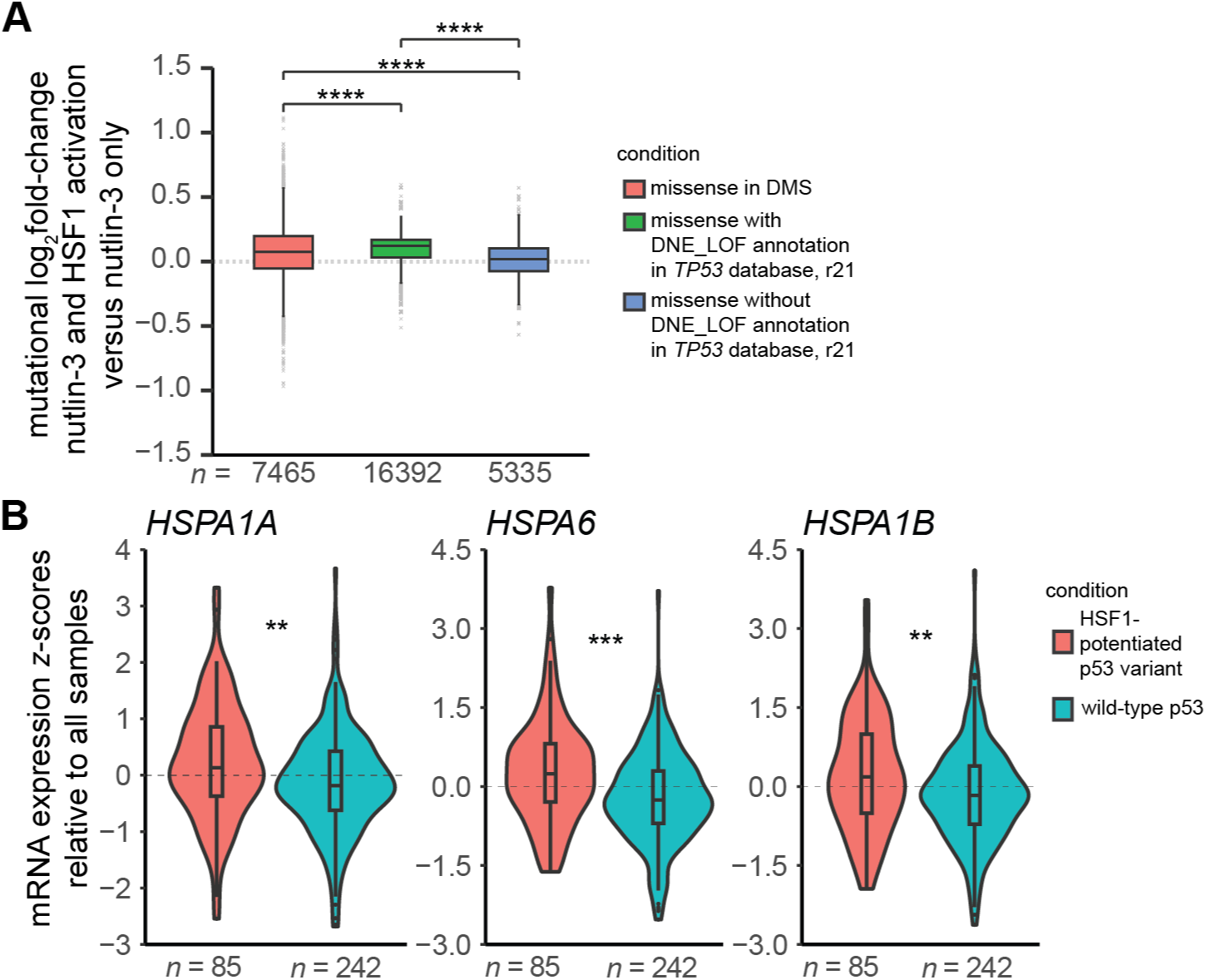
HSF1-potentiated p53 variants are enriched in patients. (**A**) Box plot comparing the effects of HSF1 in a nutlin-3 environment (mutational log_2_ fold-change nutlin-3 and HSF1 activation versus nutlin-3 only) of missense mutations found in the DMS experiment (*orange*) compared to missense mutations with (*green*) and without (*blue*) the dominant-negative loss of function (DNE_LOF) annotation in the TP53 database (r21 release). Outliers are represented as *grey* crosses. Statistical significance was calculated using a Wilcoxon-signed rank test with **** representing a *p-*value <0.0001. (**B**) Violin plots depicting mRNA expression of sentinel genes downstream of HSF1 (*HSPA1A*, *HSPA1B* and *HSPA6*) in patients expressing wild-type p53 or an HSF1-potentiated p53 variant. Statistical significance was calculated using a Wilcoxon sign rank test with ** and *** representing adjusted two-tail *p*-values <0.01 and <0.001, respectively.

We asked whether HSF1 activity levels might be higher in patients who carry an HSF1-potentiated p53 substitution. We defined “HSF1-potentiated variants” as p53 variants where the mutational log_2_ fold-change of HSF1 and nutlin-3 versus nutlin-3 alone was >0.05. Using the TCGA PanCancer Atlas of lung adenocarcinoma available in cBioPortal^85–87^, we found that expression of sentinel genes that report on HSF1 activity in patients with wild-type p53 was significantly lower than HSF1 sentinel gene activity observed in patients carrying a p53 variant potentiated by HSF1 activity in our DMS studies (**Figure 7B**). Together, these observations support the notion that HSF1 is a fundamental force shaping p53 mutational landscapes in human cancers.

## DISCUSSION

This study shows that HSF1 activity increases the fitness of p53 dominant-negative variants that can drive cancer. Moreover, our analyses of p53’s structured DNA-binding domain show that the p53 mutational fitness-enhancing impact of HSF1 activation is strongly biased towards supporting the emergence of thermodynamically destabilizing substitutions located within buried protein regions. In particular, HSF1 upregulation improves the fitness of non-conservative substitutions where buried nonpolar amino acids are replaced by polar or charged amino acids. This effect is observed across multiple cancer models. RNA-seq analysis shows that HSF1-activated genes are enriched in p53-interacting chaperones, supporting the hypothesis that HSF1 directly impacts the fitness of p53 variants through modulation of folding, aggregation, degradation, or stability of mutant p53 via HSF1-regulated chaperones. Visualization of a destabilizing p53 variant within cells under basal conditions, HSF1 activation, and HSP90 inhibition is consistent with the notion that HSF1 activation reduces aggregation of otherwise unstable p53 variants. Thus, our data are consistent with the notion that HSF1 over-activation may overcome wild-type p53 suppression to restore the dominant-negative function of mutant p53, potentially amplifying the feed-forward circuit of HSF1 activation and continued oncogenic p53 stabilization. Finally, using human cancer patient data, we find that these phenomena may be relevant in a clinical setting.

We show in this work that HSF1 can directly shape the mutational space accessible for malignant transformation. These results have several interesting implications. While the literature has clearly highlighted HSF1 activation as an oncogenic helper^7, 88, 89^, it is unclear whether the commonly observed increase in HSF1 expression and activity occurs prior to the onset of tumorigenesis, or whether constitutive HSF1 activation is a consequence of proteome instability arising from accumulated mutations and genomic damage. Our results suggest that HSF1 activation can facilitate early tumorigenic events by increasing the fitness of some oncogenic driver mutations. Second, resistance to chemo-therapeutic agents is frequently driven by mutations within the targeted proteins. While both HSF1 upregulation as well as downstream chaperones have been identified as facilitators of chemo-resistance, the mechanism has primarily been associated with alterations in metabolic or autophagy pathways. Further studies on HSF1 targets that interact with p53 should be pursued to better understand the interplay between chaperone networks and oncogenesis ^18, 19, 90, 91^. Our results suggest that HSF1 activity may facilitate chemo-resistance by tuning the accessibility of resistance mutations. Given the recent therapeutic interest in inhibitors of both HSF1 itself and HSF1-regulated chaperones such as HSP90 and HSP70^22, 23, 52, 59^, these results indicate that such inhibitors may be particularly effective in combination therapy to prolong effective outcomes by reducing the emergence of resistant tumor populations^92^.

The capacity of HSF1 or other types of proteostasis network upregulation to tune the stability, expression levels, folding, and/or aggregation of oncoproteins could have tremendous implications for disease treatment^93^. Variants with enhanced stability may not be able to be degraded, and thus, unable to be efficiently presented to the immune system by MHC-1 proteins. In accordance with our findings and published work in this field, we reason that downregulation of proteostasis networks may enable a more robust immune response to tumor presenting antigens^94^.

We emphasize that the impacts of proteostasis network modulation on oncoprotein mutational spectra are likely to extend far beyond just p53. This study should motivate efforts to more fully understand how HSF1 shapes the mutational spectra of additional oncoproteins and chemotherapy targets.

## LIMITATIONS OF STUDY

While we observed a loss of fitness of HSF1-potentiated p53 variants with chaperone inhibition, HSP90 and HSP70 inhibition can also induce compensatory HSF1 activation^59^. Untangling the relative effects of chaperone activity versus HSF1 activity is challenging. Considering these competing phenomena would require a comprehensive understanding of the effects of individual chaperones on individual p53 substitutions. Additionally, while we focused our follow-up experiments on sentinel HSF1-targeted genes (HSP70 and HSP90), other HSF1-induced p53 interactors may also play a role in the relative fitness of p53 variants. These effects could include direct interactions with p53 or secondary effects due to remodeling of the proteostasis environment and requires targeted mechanistic work on individual p53 variants. Overall, while the results we report implicate chaperone inhibition or HSF1 inhibition as potential therapeutic targets for cancers harboring a p53 variants, inhibitor concentrations will need to be tuned or used in combination with other therapeutics. Further studies directly testing this concept are needed to draw a robust conclusion.

A second caveat to consider is that the DMS library only contains single amino acid substitutions in p53 and the mutant library was transduced via lentiviral overexpression. Large deletions, multiple missense mutations, or copy number variations, which can all impact p53 biology and play a role in tumorigenesis, were not considered. HSF1 may play a role in such cases due to the proteostasis imbalances, and this phenomenon should be further studied.

Finally, while our study provides evidence for roles of HSF1 activity in oncogenic mutational spectra across multiple cell lines, we are still limited by the model system. Our work aimed to intentionally induce HSF1 not via stress, however chronic stress in the tumor microenvironment and subsequent HSF1 activation could lead to differing outcomes. The work here should motivate related future *in vivo* studies.

## RESOURCE AVAILABILITY

### Lead contact

Requests for further information or requests for resources and reagents should be directed to, and will be fulfilled by, the lead contact, Matthew D. Shoulders.

### Materials availability

Plasmids and cell lines generated from this study are available from the lead contact upon request and with a completed material transfer agreement.

### Data and code availability

Sequencing data have been deposited at SRA as SRA: PRJNA1178283 and are publicly available as of the date of publication. RNA-seq data from this paper (**Tables S1** and **S7**) have been deposited at GEO as GEO:GSE304320 and are publicly available as of the date of publication.

## ACKNOWLEDGEMENTS

This work was supported by the National Institutes of Health (1R35GM136354 to M.D.S.), (1R01AI168166 to Y.-S.L), MIT HEALS (to M.D.S.), and by an American Cancer Society–Ellison Foundation Research Scholar Award (to M.D.S.). Additional support was provided by Koch Institute Support (core) under NIH (core) grant P30-CA14051 from the NIH/NCI and by the MIT CEHS core via the NIH/NIEHS (Grant P30-ES002109). Work in the Sánchez-Rivera laboratory is supported by the Howard Hughes Medical Institute (Hanna Gray Fellowship, GT15656), V Foundation for Cancer Research (V2022-028), NCI 1P01CA291694-01A1, Virginia and D.K. Ludwig Fund for Cancer Research, MIT HEALS Initiative, Koch Institute Frontier Research Program, Casey and Family Foundation Cancer Research Fund, Michael (1957) and Inara Erdei Fund, MIT Research Support Committee, Upstage Lung Cancer Foundation, and a Traditional Project Award from the Bridge Project, a partnership between the Koch Institute for Integrative Cancer Research at MIT and the Dana-Farber/Harvard Cancer Center. P.R. was supported by a National Science Foundation GRFP Award.

## AUTHOR CONTRIBUTIONS

S.H, R.M.S., K.E.L, T.H., A.O.G., Y.-S.L., and M.D.S. designed experiments. S.H, K.E.L, R.M.S., and T.H. conducted experiments. R.M.S. and M.D.S. conceived the project. A.O.G. and W.C.H. provided key reagents. All authors analyzed results and contributed to writing and/or editing the manuscript. Y.-S.L. and M.D.S. supervised the research.

## STAR Methods

### METHOD DETAILS

#### Cell culture

A549 cells were a kind gift from the Prof. William Hahn Lab at Harvard Medical School. U2OS cells were purchased from ATCC. Cells were grown in DMEM medium (Corning), supplemented with 10% heat-inactivated fetal bovine serum (FBS, Cellgro) and 1% penicillin/streptomycin/glutamine (Cellgro) at 37 °C with 5% CO_2_(g).

#### Plasmids

To create stable A549^cHSF1^ and U2OS^cHSF1^ cell lines, cHSF1^59^ was cloned into the pINDUCER20 lentiviral vector (AddGene #44012) using Gateway cloning. pINDUCER20 expresses both a gene of interest under a tetracycline responsive TRE2 promoter as well as a tetracycline activator (rtTA3) under a constitutive promoter, enabling inducible regulation of a gene of interest following a single lentiviral transduction. The *TP53* library was expressed in a modified lentiviral pMT_BRD025 vector (AddGene #113569)^67^. To generate cell lines co-expressing p53 variants, fluorescent markers down-stream of p53 and a t2a ribosome skip sequence were cloned into LX313-TP53-WT (Addgene #118014) plasmid using Gibson assembly (NEB E2621S). Site-directed mutagenesis (NEB E0554S) was used to generate point mutations in the p53 gene.

#### Lentivirus production

LentiX cells (Takara Bio), cultured as described above, were co-transfected with the structural plasmids necessary for virus production (psPAX2 and pMDM2.G from AddGene) along with the lentiviral vectors for either pINDUCER20.cHSF1, LX313-TP53-WT or variants, or the *TP53* mutational library. Cells were transfected using TransIT-Lenti (Mirus) for 24 h, after which the media was removed and replaced with fresh media. Media containing viral particles was collected at 48 h and cell debris was removed by centrifugation at 500 × *g* for 10 min. Viral supernatant was either concentrated using lentivirus precipitation solution (PEG-IT SBI) or aliquoted and stored at −80 °C until use. To measure the titer of the *TP53* library lentivirus, A549 cells were infected with serially diluted virus in 96-well plates. The infected cells were then selected in puromycin (Gibco) for 48 h and surviving cells were quantified using resazurin (Sigma).

#### Lentiviral titer assay

A549^cHSF1^ cells were seeded in 96-well plates (Corning) at a density of 3 × 10^5^ cells/well in DMEM medium. The following day media was removed and replaced with viral media containing polybrene at a final concentration of 8 μ/mL. After a 96-h incubation, media was removed and replaced with 100 μL of DMEM containing 0.01 mg/mL resazurin sodium salt (Sigma). After 2 h of incubation, resorufin fluorescence (excitation 530 nm; emission 590 nm) was quantified using a Take-3 plate reader (BioTeK). Experiments were conducted in biological triplicate. Viral titer in transducing units per mL (TU/mL) was calculated as: [(number of cells plated) x (fraction of surviving cells)] / (volume of virus). The average of the calculated TU/mL over the linear range of the assay was used for subsequent calculation of appropriate multiplicity of infection.

#### Stable cell line engineering

For the construction of A549^cHSF1^ cells, A549 cells were transduced with lentivirus co-encoding a G418-resistance gene and rtTA3 alongside cHSF1 in the presence of 2 μg/mL polybrene (Sigma-Aldrich). Heterostable cell lines were then selected using 1 mg/mL G418 (Enzo Life Sciences). Clonal populations were screened based on functional testing of the cHSF1 construct using real-time polymerase chain reaction (RT-PCR; described below) with or without 1 μg/mL dox (Alfa Aesar). For the construction of U2OS^cHSF1^ cells, U2OS cells were transduced with lentivirus co-encoding a G418-resistance gene and rtTA3 alongside cHSF1 in the presence of 2 μg/mL polybrene (Sigma-Aldrich). Heterostable cell lines were then selected using 150 μg/mL G418 (Enzo Life Sciences) and functional testing of the cHSF1 construct was conducted using RT-PCR with or without 0.01 μg/mL dox.

#### Resazurin viability assay

A549^cHSF1^ cells were seeded in 96-well plates (Corning) at a density of 3 × 10^5^ cells/well in DMEM medium and then treated with 0.1% DMSO, 1 μg/mL dox, 2.5 μM nutlin-3 (Cayman Chemical Company), or 1 μg/mL dox and 2.5 μM nutlin-3. 48 h post-treatment, media was removed and replaced with 100 μL of DMEM containing 0.01 mg/mL resazurin sodium salt (Sigma). After 2 h of incubation, resorufin fluorescence (excitation 530 nm; emission 590 nm) was quantified using a Take-3 plate reader (BioTeK). Experiments were conducted in biological triplicate.

#### RT-PCR

A549^cHSF1^ cells were treated with 1 μg/mL dox for 24 h for assessment of cHSF1 construct function, while a 6 h treatment with 500 nM STA-9090 (MedChem Express) was used as a positive control for HSR activation. RNA was extracted using the EZNA Total RNA Kit I (Omega). qRT-PCR reactions were performed on cDNA prepared from 1000 ng of total cellular RNA using the High-Capacity cDNA Reverse Transcription Kit (Applied Biosystems). The Fast Start Universal SYBR Green Master Mix (Roche) and appropriate primers purchased Sigma were used for amplifications (6 min at 95 °C then 45 cycles of 10 s at 95 °C, 30 s at 60 °C) in a Light Cycler 480 II Real-Time PCR machine. The primers used for *DNAJB1* were 5′-TGTGTGGCTGCACAGTGAAC-3′ (forward) and 5′-AC-GTTTCTCGGGTGTTTTGG-3′ (reverse), primers for *HSPA1A* were 5′-GGAGGCGGAGAAGTACA-3′ (forward) and 5′-GCTGATGATGGGGTTACA-3′ (reverse), primers for *RPLP2* were 5′-CCATTCAGCTCACTGATAACCTTG-3′ (forward) and 5′-CGTCGCCTCCTACCTGCT-3′ (reverse), primers for *CDKN1A* were 5′-CGACTGTGATGCGCTAATGG-3′ (forward) and 5′-CCGTGG-GAAGGTAGAGCTTG-3′ (reverse), and primers for *BAX* were 5′-CAGCAAACTGGTGCTCAAGG-3′ (forward) and 5′-TCCTGGAGACAGGACATCA-3′ (reverse). Transcripts were normalized to the house-keeping genes *RPLP2*. All measurements were performed in technical triplicate. Data were analyzed using the LightCycler® 480 Software, Version 1.5 (Roche) and data are reported as the mean ±95% confidence intervals.

#### RNA-Seq

A549^cHSF1^ cells were seeded at 7.5 x 10^4^ cells/well in a 12-well plate in DMEM media. Cells were then treated with either 0.01 % DMSO or 1 μg/mL dox for 24 h. Cellular RNA was harvested using the RNeasy Plus Mini Kit with QIAshredder homogenization columns (Qiagen). RNA samples were quantified using an Advanced Analytical Fragment Analyzer. The initial steps were performed on a Tecan EVO150.10 ng of total RNA was used for library preparation. 3′DGE-custom primers 3V6NEXT-bmc#1-24 were added to a final concentration of 1 μM. (5’-/5Biosg/ACACTCTTTCCCTACACGAC-GCTCTTCCGATCT [BC6]N10T30VN-3′ where 5Biosg = 5′ biotin, [BC6] = 6bp barcode specific to each sample/well, N10 = Unique Molecular Identifiers, Integrated DNA technologies) were used to generate two subpools of 24 samples each^95, 96^. After addition of the oligonucleotides, Maxima H Minus RT was added per the manufacturer’s recommendations with the template-switching oligo 5V6NEXT (10 μM, [5V6NEXT: 5′-iCiGiCACACTCTTTCCCTACACGACGCrGrGrG-3′ where iC: iso-dC, iG: iso-dG, rG: RNA G]), followed by incubation at 42 °C for 90 min and inactivation at 80 °C for 10 min. Following the template switching reaction, cDNA from 24 wells containing unique well identifiers were pooled together and cleaned using RNA Ampure beads at 1.0×. cDNA was eluted with 17 μL of water followed by digestion with Exonuclease I at 37 °C for 30 min, and inactivation at 80 °C for 20 min. Second strand synthesis and PCR amplification was done by adding the Advantage 2 Polymerase Mix (Clontech) and the SINGV6 primer (10 pM, Integrated DNA Technologies 5′-/5Biosg/ACACTCTTTCCCTACACGACGC-3′) directly to half of the exonuclease reaction volume. Eight cycles of PCR were performed, followed by clean-up using regular SPRI beads at 0.6×, and elution with 20 μL of Resuspension Buffer (Illumina). Successful amplification of cDNA was confirmed using the Fragment Analyzer. Illumina libraries were then produced using Nextera FLEX tagmentation substituting P5NEXTPT5-bmc primer (25 μM), Integrated DNA Technologies, (5′-AATGATACGGCGACCACCGAGATCTACACTCTTTCCCTACACGAC-GCTCTTCCG*A*T*C*T*-3′ where * = phosphorothioate bonds) in place of the normal N500 primer. Final libraries were cleaned using SPRI beads at 0.7× and quantified using the Fragment Analyzer and qPCR before being loaded for paired-end sequencing using the Illumina NextSeq500 in paired-end mode (26/50 nt reads).

Analyses were performed using previously described tools and methods^97^. Reads were aligned against hg19 (Feb., 2009) using bwa mem v. 0.7.12-r1039 [RRID:SCR_010910] with flags –t 16 –f, and mapping rates, fraction of multiply-mapping reads, number of unique 20-mers at the 5’ end of the reads, insert size distributions, and fraction of ribosomal RNAs were calculated using bedtools v. 2.25.0 [RRID:SCR_006646]^98^. In addition, each resulting bam file was randomly down-sampled to a million reads, which were aligned against hg19, and read density across genomic features were estimated for RNA-Seq-specific quality control metrics. For mapping and quantitation, reads were scored against GRCh38/ENSEMBL 101 annotation using Salmon v.1.3 with flags quant -p 8 -l ISR –validate-Mappings^99^. The resulting quant.sf files were imported into the R statistical environment using the tximport library (tximport function, option “salmon”), and gene-level counts and transcript per-milllion (TPM) estimates were calculated for protein-coding genes. Samples were clustered based on genes with aver-age log2 TPM >0.1 across all samples (n=6320 genes) based on complete linkage clustering of the Co-sine correlation among samples. Samples with similarity score <0.94, which were clear outliers from the rest, were excluded from further analysis (n=5).

Differential expression was also analyzed in the R statistical environment (R v.3.5.1) using Bioconductor’s DESeq2 package on the protein-coding genes only [RRID:SCR_000154]^100^. Dataset parameters were estimated using the estimateSizeFactors(), and estimateDispersions() functions; read counts across conditions were modeled based on a negative binomial distribution, and a Wald test was used to test for differential expression (nbinomWaldtest(), all packaged into the DESeq() function), using the treatment type as a contrast. Shrunken log_2_ fold-changes were calculated using the lfcShrink function, based on a normal shrinkage estimator^100^. Fold-changes and *p*-values were reported for each protein-coding gene. Upregulation was defined as a change in expression level >1.5-fold relative to the basal environment with a non-adjusted *p*-value < 10^−5^. Gene ontology analyses were performed using the online DAVID server, according to tools and methods presented by Huang and co-workers^97^.

#### Gene set enrichment analysis (GSEA)

Differential expression results from DESeq2 were retrieved, and the “stat” column was used to pre-rank genes for GSEA analysis. These “stat” values reflect the Wald’s test performed on read counts as modeled by DESeq2 using the negative binomial distribution. Genes that were not expressed were excluded from the analysis. GSEA (linux desktop version, v4.1)^101, 102^ was run in the pre-ranked mode against MSigDB 7.4 C5 (Gene Ontology) set, and ENSEMBL IDs were collapsed to gene symbols using the Human_ENSEMBL_Gene_ID_MSigDB.v7.4.chip (resulting in 12706 unique genes for par and 12141 for sg4, respectively). In addition, a weighted scoring scheme, meandiv normalization, and cutoffs on MSigDB signatures sizes (between 5 and 2000 genes, resulting in 8496 gene sets retained) were applied and 5000 permutations were run for *p*-value estimation.

#### Generating A549^cHSF1^(p53-Lib) cells

A549^cHSF1^ cells were infected with titered p53 library lentivirus at a multiplicity of infection of 0.25 (4 × 10^7^ cells mixed with 1 × 10^7^ lentiviral particles) in the presence of 8 μg/mL polybrene. Following transduction, cells were selected with 2 μg/mL puromycin (Gibco).

#### Deep mutational scanning

A549^cHSF1^(p53-Lib) cells were seeded in 15 cm tissue culture plates at a density of 3 × 10^6^ cells/plate. In order to maintain library diversity throughout selection, three plates were used per treatment for a total of 9 × 10^6^ cells. Cells were treated with 0.01% DMSO, 1 μg/mL dox, 0.01% DMSO and 2.5 μM nutlin-3, or 1 μg/mL dox and 2.5 μM nutlin-3. Cells were trypsinized, counted, and re-seeded in three plates each at 3 × 10^6^ cells/plate every 3 d. Following 12 d of treatment, cell pellets were harvested by centrifugation at 1,000 rpm for 5 min. Aliquots of 9 × 10^6^ cells were snap-frozen in liquid N_2_(g) in Eppendorf tubes and stored at –80 °C for subsequent DNA extraction. The deep mutational scanning experiment was repeated independently for a total of three biological replicates from the same p53-Lib cell line.

To prepare samples for Illumina sequencing, genomic DNA was purified from aliquots of frozen cells using the QIAamp Blood Midi Kit (Qiagen) and final DNA concentration was determined using a Qubit (Fischer). PCR amplicons of p53 were prepared using 2.0 μg of genomic DNA over 25 cycles and with Herculase II as the DNA polymerase (Agilent). The primers used were 5’ ATTCTCCTTGGAATTT-GCCCTT 3’ and 5’ CATAGCGTAAAAGGAGCAACA 3’. Twelve PCR reactions were performed per sample, and the reactions were pooled and cleaned up using a PCR clean-up kit (Omega). The p53 amplicons were further gel-purified using a Pippin prep system (Sage Science) prior to library preparation via Nextera Flex. The resulting libraries were quantified using the Fragment Analyzer before they were pooled and sequenced on an Illumina NovaSeq with 2 × 150 bp paired-end reads.

#### Deep mutational scanning data analysis

The software ORFCall v1.0 [https://github.com/broadinsti-tute/ORFCall/releases/tag/v1.0] was used with flags -p -Q 30 to align the deep-sequencing reads against the *TP53* wild-type sequence and count the number of times each codon mutation was observed in each selection condition. The mutational fold-change for each variant was calculated by normalizing raw read counts to the total read count at each position. Next, the log_2_ fold-change in selection versus mock conditions was calculated by taking the log of mutational fold-change in the selection condition normalized to the mutational fold-change in the mock condition. Mutational fitness in each condition was then determined by averaging the log_2_ fold-change from three biological replicates. RSA was calculated using the software DSSP on chain A of the p53 DNA-binding domain crystal structure (PDBID 2OCJ)^103, 104^. DSSP calculates the solvent-accessible surface area of the monomer (ASA) and the RSA is calculated by dividing the ASA by the total theoretical solvent accessibility area^105^. Sites were classified as buried if the RSA was <0.2 and exposed if the RSA was >0.2.

#### Rosetta analysis

The calculations for ΔΔ*G* of protein stability upon substitution were performed using the cartesian_ddg application in Rosetta version 3.13^80^. The crystal structure of the DNA-binding domain of p53 (PDB ID: 2OCJ, chain A) was used as the initial structure for the ΔΔ*G* calculations^104^. The initial p53 structure was relaxed using the Rosetta FastRelax application to generate a total of 20 relaxed decoys. The Rosetta FastRelax application performed five cycles of side-chain repacking and energy minimization with the Rosetta energy function ref2015_cart^80, 106–108^. The lowest energy structure of the 20 decoys was used as the wild-type structure for the cartesian_ddg calculation. In the carte-sian_ddg calculation, the target residue was substituted with each of the 20 natural amino acids, and any neighboring residues within a 9-Å radius were repacked and energy-minimized using the ref2015_cart energy function. This calculation process was performed five times to generate five energy scores for the mutant and for the wild-type. The average wild-type scores were subtracted from the average mutant scores to calculate the ΔΔ*G* values. The ΔΔ*G* values were then scaled by a factor of 0.34; this scale factor was previously calculated by fitting Rosetta-predicted ΔΔ*G* values to experimental ΔΔ*G* values in units of kcal/mol, and is used here to better relate predicted ΔΔ*G* values to experimental values^80^.

#### Flow cytometry-based pairwise competition assay

A549^cHSF1^ and U2OS^cHSF1^ cells were transduced with concentrated lentivirus co-encoding a hygromycin-resistance gene, wild-type or a variant of p53, and a fluorescent protein. Transduced cells were then selected using 1 mg/mL of hygromycin-B for A549^cHSF1^ cells or 100 μg/mL of hygromycin-B for U2OS^cHSF1^ cells for 5 d. 25,000 wildtype + BFP cells with 25,000 variant + GFP cells were plated in one well of a 12-well plate, and the remaining cells were used for flow analysis to quantify the pre-competition mix. Cells were then treated with DMSO as vehicle or dox (1 μg/mL for A549^cHSF1^ cells and 0.01 μg/mL for U2OS^cHSF1^ cells), 2.5 μM nutlin-3, 500 nM STA-9090 (MedChem Express), 5 μM VER-155008 (Med-Chem Express) 24 h after plating. 48–96 h post-treatment, the GFP+ fraction of the cells was evaluated via flow cytometry. Fluorescence-based measurements for the validation of p53 variants were performed using the BD LSRFortessa Cell Analyzer in tube or plate reader format, using the BD FACSDiva v.3.0 software for data collection. Log_2_ fold-change of the GFP+ cell fraction was calculated by first normalizing to the pre-competition mix and then to either the average of the vehicle-treated or nutlin-3-alone-treated cells.

#### Immunohistochemistry and confocal microscopy

A549^cHSF1^ cells expressing p53 variants and a fluorescent protein were plated in a glass bottom 6-well plate (Cellvis). Cells were treated with drugs 24 h after plating. Cells were washed with PBS then fixed with 4% paraformaldehyde (Electron Microscopy Sciences) for 25 min at RT. Samples were treated with 0.1% Triton X-100 (Thermo Scientific) for 30 min to permeabilize the cells. Samples were then incubated in a solution of 5% BSA (Gibco) in PBS at RT for 30 min to block non-specific antibody binding. Cells were labeled with anti-p53 antibody (Santa-Cruz, sc-126, DO-1) in 1% BSA for 1 h at RT. Samples were washed 3× with PBS. Samples were incubated with Alexa Fluor 568-conjugated anti-mouse (Invitrogen) in 1% BSA for 1 h at RT. Samples were stained with DAPI (Thermo Fisher). Images were acquired at the Swanson Biotechnology Center at the Koch Institute on an Evident FV4000 with a 100× oil-immersion objective (UPLSAPO100XS). The cellSens FV software was used for image acquisition. The excitation lasers to capture the images were 405, 488 and 561 nm. Image processing was performed using ImageJ.

#### Analysis of TCGA data

Human data from the TCGA lung adenocarcinoma study, particularly *TP53* mutations and *HSPA1A, HSPA6,* and *HSPA1B* mRNA expression data, were downloaded from cBioPortal. Study name: Lung adenocarcinoma TCGA, PanCancer Atlas, 507 total samples. We split the data into patients expressing wild-type and variants of p53. We then selected patients with variants of p53 that were found to have a mutational log_2_ fold-change nutlin-3 and HSF1 versus nutlin-3 alone >0.05 in our DMS experiment.

### QUANTIFICATION AND STATISTICAL ANALYSIS

#### Statistical analyses

All experiments were performed in biological triplicate. Statistical analyses were done in Prisim GraphPad, Jupyter Notebook, and R. Site and mutational log_2_ fold-change (**Figure 2**) were calculated using a Wilcoxon signed-rank test. Statistical significance in mutational log_2_ fold-change in missense mutations and buried versus exposed sites (**Figures 4B**, **S2C**, **S2D**, **S6B**, and **S6D**) were calculated using a Welch’s *t*-test for independent samples with Bonferroni correction, and significance from null was determined using a Wilcoxon signed-rank test. As a non-parametric test, the Wilcoxon’s rank-sum test is particularly adept at assessing whether the sample comes from a population that is symmetrically distributed around a center (in our case, 0) without assuming normality of the distribution. All correlations were determined by calculating Pearson correlation coefficients using a two-tailed test. Statistical significance between exposed and buried sites (**Figure 4C**) and mRNA expression change between wild-type and HSF1 potentiated variants (**Figure 7C**) were calculated using a Wilcoxon-signed rank test. The statistical significance between solvent RSA classes or mutation types within a solvent accessibility class (**Figures S6C** and **S6D**) were evaluated using ANOVA, while comparisons between select conditions were calculated using Welch’s *t*-test for independent samples with Bonferroni correction. We applied pairwise Welch-corrected *t*-tests in case of potential unequal variance among samples. Additionally, to control for type I errors, we applied a stringent Bonferroni correction to the resulting pairwise-comparison *p*-values. Statistical significance between stabilizing and destabilizing mutations based on experimental measurements (**Figure S6F**) as well as for significance between buried and exposed regions (**Figure 5B**) were calculated using a Welch’s *t-*test for independent variables. ANOVA was also used to calculate statistical significance between RSA class and amino acid changes and ΔΔ*G* (**Figure 5E**) and between missense mutations in the DMS experiment and *TP53* database (**Figures 7A** and **7B**). Statistical significance for flow cytometry-based competition assays was calculated using Fisher’s exact test for counts data (biological replicates were summed prior to calculating significance) (**Figures 5E–H, 6D**, and **6E**). Complete outcomes of Fisher’s exact tests with odds ratio and 95% confidence interval are reported in **Table S6**.

## Supplemental Excel table titles

**Table S1:** RNA-Seq differential expression analysis of A549^cHSF1^ cells and GSEA. Related to Figure 1.

**Table S2:** APID p53 interactors. Related to Figure 1.

**Table S3:** DMS experiment full data (*TP53* library coverage. Mutational log_2_ fold-change values. Site log_2_ fold-change values.) Related to Figure 2.

**Table S4:** Surface accessible solvent area. Related to Figure 4.

**Table S5:** Complete Rosetta ΔΔ*G* analysis. Related to Figure 4.

**Table S6:** Pairwise competition data and statistics. Related to Figures 5 and 6.

**Table S7**: RNA-seq differential expression analysis of A549^cHSF1^, A549^dn-cHSF1^ and A549 cells treated with dox.

