## Supplemental figures for "Dominant-negative *TP53* mutations potentiated by the HSF1-regulated proteostasis network"


| Page | Content |
| --- | --- |
| S1 | Table of Contents |
| S2 | List of Supporting Tables |
| S3–11 | Supporting Figures and Captions |

**Table S1:** RNA-Seq differential expression analysis of A549$^{cHSF1}$ cells and GSEA. Related to Figure 1.
**Table S2:** APID p53 interactors. Related to Figure 1.
**Table S3:** DMS experiment full data (*TP53* library coverage. Mutational $\log_2$ fold-change values. Site $\log_2$ fold-change values.) Related to Figure 2.
**Table S4:** Surface accessible solvent area. Related to Figure 4.
**Table S5:** Complete Rosetta $\Delta\Delta G$ analysis. Related to Figure 4.
**Table S6:** Pairwise competition data and statistics. Related to Figures 5 and 6.
**Table S7**: RNA-seq differential expression analysis of A549$^{cHSF1}$, A549$^{dn\text{-}cHSF1}$ and A549 cells treated with dox.

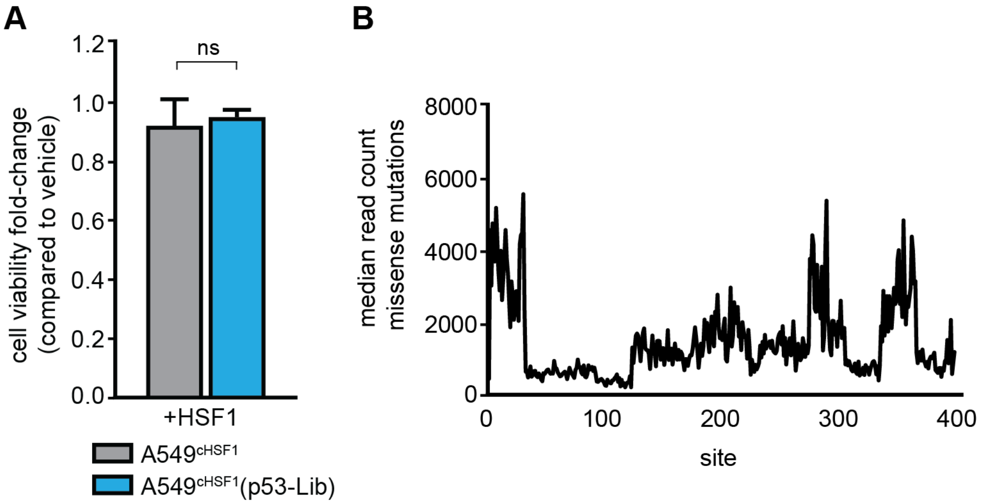


**Figure S1: Validation of A549$^{cHSF1}$ cell lines expressing the *TP53* mutational library. Related to Figure 1**

(**A**) Induction of HSF1 did not significantly impair the viability of A549$^{cHSF1}$ cells that either expressed or did not express the p53 library, as measured by a resazurin assay. The average cell viability fold-change of biological triplicates is plotted with error bars representing the standard deviation. (**B**) Median read count for each codon plotted against the amino acid site number. Data for library coverage are provided in **Table S1**.

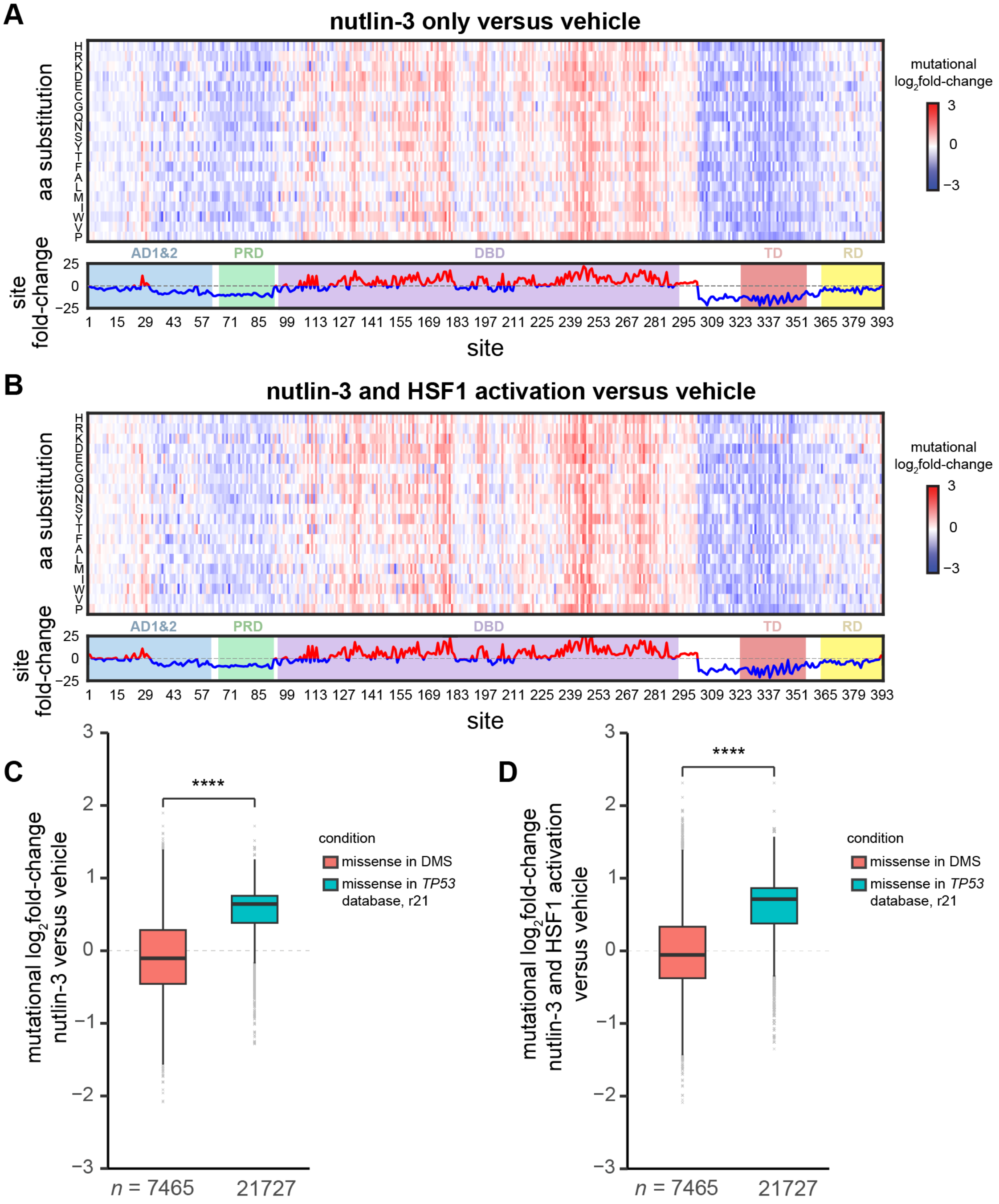


**Figure S2: Enrichment of cancer-associated, dominant negative p53 variants upon nutlin-3 selection. Related to Figure 2.**

Heat map of the *TP53* mutational $\log_2$ fold-change averaged over three biological replicates for each amino acid substitution for (**A**) nutlin-3 treatment of A549$^{cHSF1}$(p53-Lib) in a basal proteostasis environment and (**B**) nutlin-3 treatment in a HSF1-activated proteostasis environment, as compared to the corresponding vehicle-treated (no nutlin-3) control samples. The sum of the mutational $\log_2$ fold-change at each site is shown below each heat map. Box plots for *TP53* missense mutations in the *TP53* database as compared to all missense mutations from the DMS experiment, shown here for (**C**) nutlin-3 selection in a basal proteostasis environment and (**D**) nutlin-3 selection in a cHSF1-activated proteostasis environment as compared to the corresponding vehicle-treated (no nutlin-3) control. Statistical significance was assessed using a Wilcoxon signed-rank test. **** represents a $p$-value $<0.0001$.

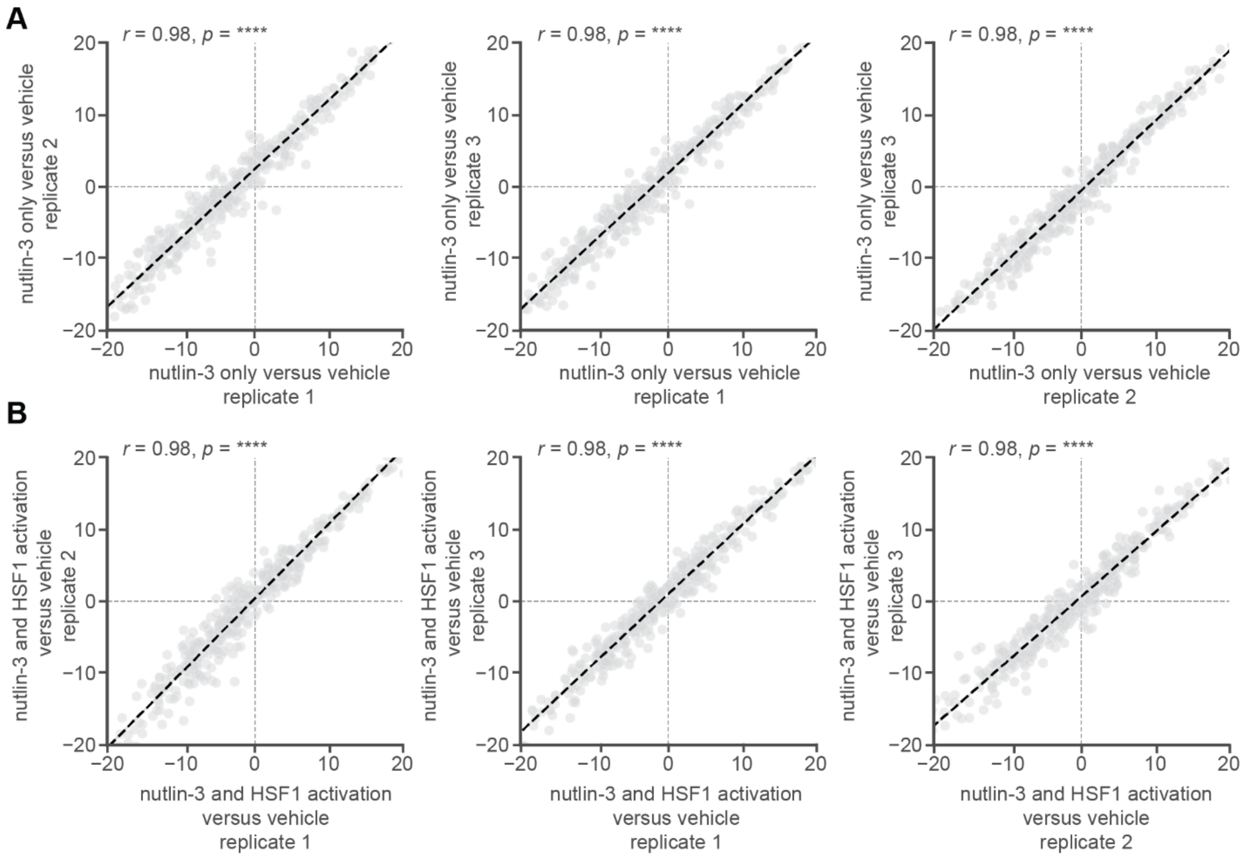


**Figure S3: Correlation between biological replicates of nutlin-3 treatment. Related to Figure 2.** Correlation between cumulative site $\log_2$ fold-change for each biological replicate for the following comparisons: (**A**) nutlin-3 versus vehicle treatment and (**B**) nutlin-3 with HSF1 activated versus vehicle treatment. Pearson correlation coefficients ***r*** as well as the corresponding *p*-values are included, with **** representing an adjusted two-tailed *p*-value <0.0001.

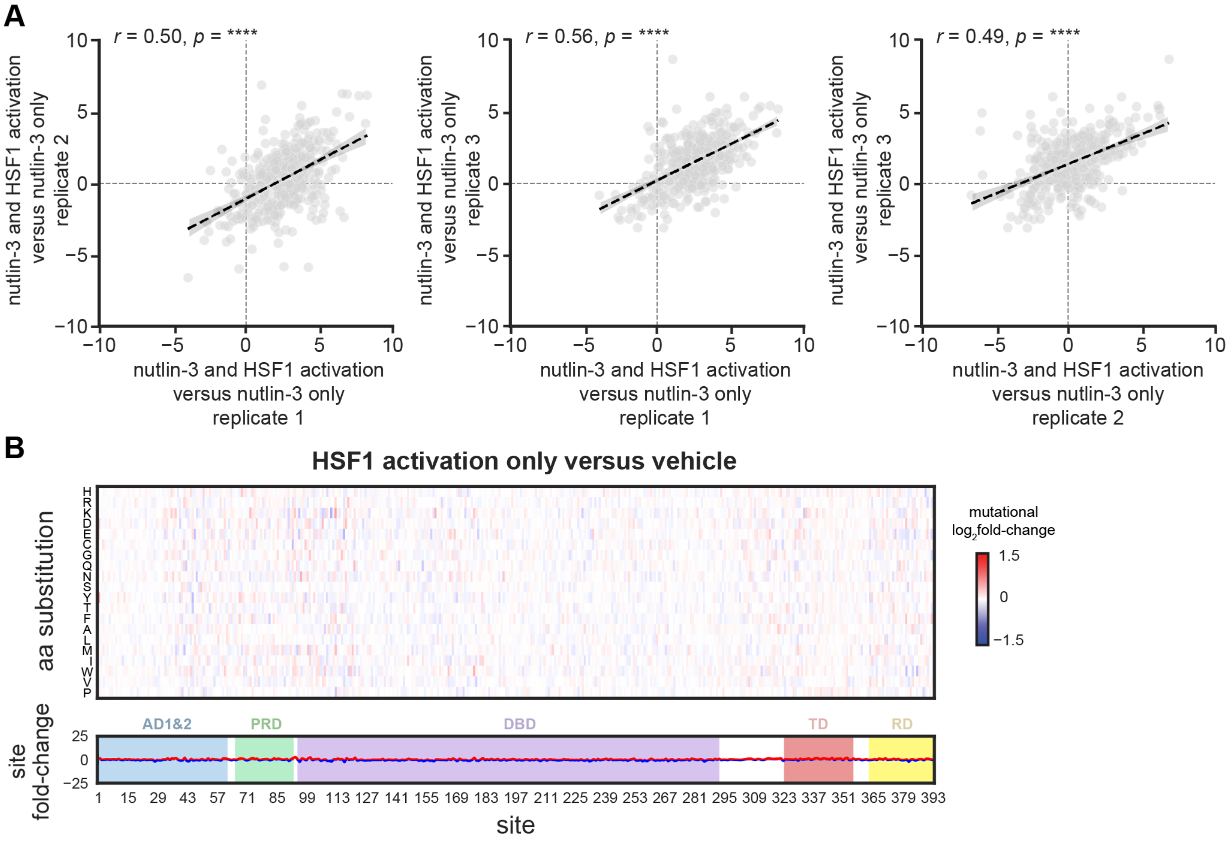


**Figure S4: Biological replicates of HSF1 activation in the context of nutlin-3 treatment and consequences of HSF1 activation in the absence of nutlin-3 treatment. Related to Figure 2.**

(**A**) Correlation between cumulative site $\log_2$ fold-change for each biological replicate for nutlin-3 treatment with cHSF1 activation versus nutlin-3 only. Pearson correlation coefficients *r* as well as the corresponding *p*-values are included, with **** representing an adjusted two-tailed *p*-value <0.0001. (**B**) Heat map of the p53 mutational frequency $\log_2$ fold-change averaged over three biological replicates for each amino acid substitution in an HSF1-activated versus basal proteostasis environment in the absence of nutlin-3 treatment. The sum of the mutational $\log_2$ fold-change at each site is shown below. Since endogenous p53 is always present, and there is no nutlin-3 to drive selection, the effects are minimal.

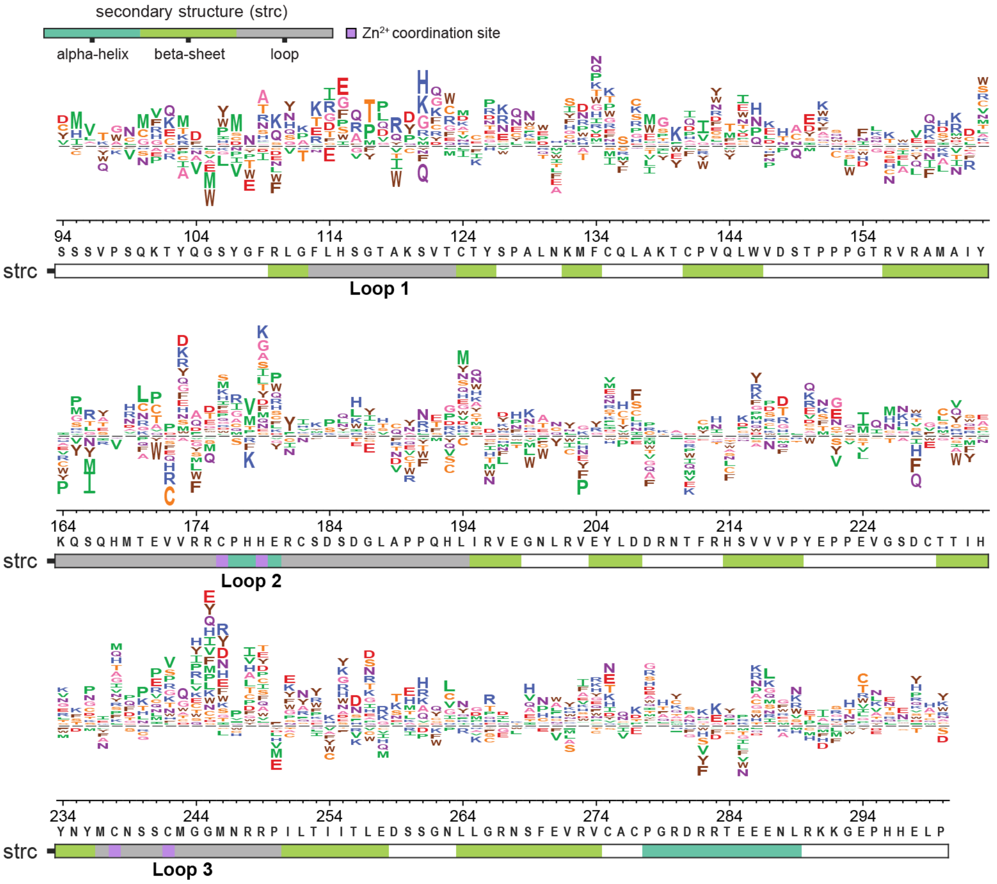


**Figure S5: Effects of HSF1 activation on p53 mutational fitness under nutlin-3 selection within the p53 DNA-binding domain. Related to Figure 3.**

Sequence logo plots for the p53 DNA-binding domain under HSF1-activated versus basal proteostasis environments during nutlin-3 selection. Alleles displaying opposing signs between biological replicates were filtered, so all data shown are for robust variants that displayed a consistent response across replicates. The color bar below the sequence logo plot indicates secondary structure elements as alpha-helical (*teal*), β-sheet (*green*) or loop (*grey*). $Zn^{2+}$ coordination sites are marked as purple. Colorization of amino acids within the logo-plot is based on the side-chain properties as follows: negatively charged (D, E; *red*), positively charged (H, K, R; *blue*), large polar (N, Q; *purple*), polar uncharged (C, S, T; *yellow*), small nonpolar (A, G; *pink*), aliphatic (I, L, M, P, V; *green*), and aromatic (F,W,Y; *brown*).

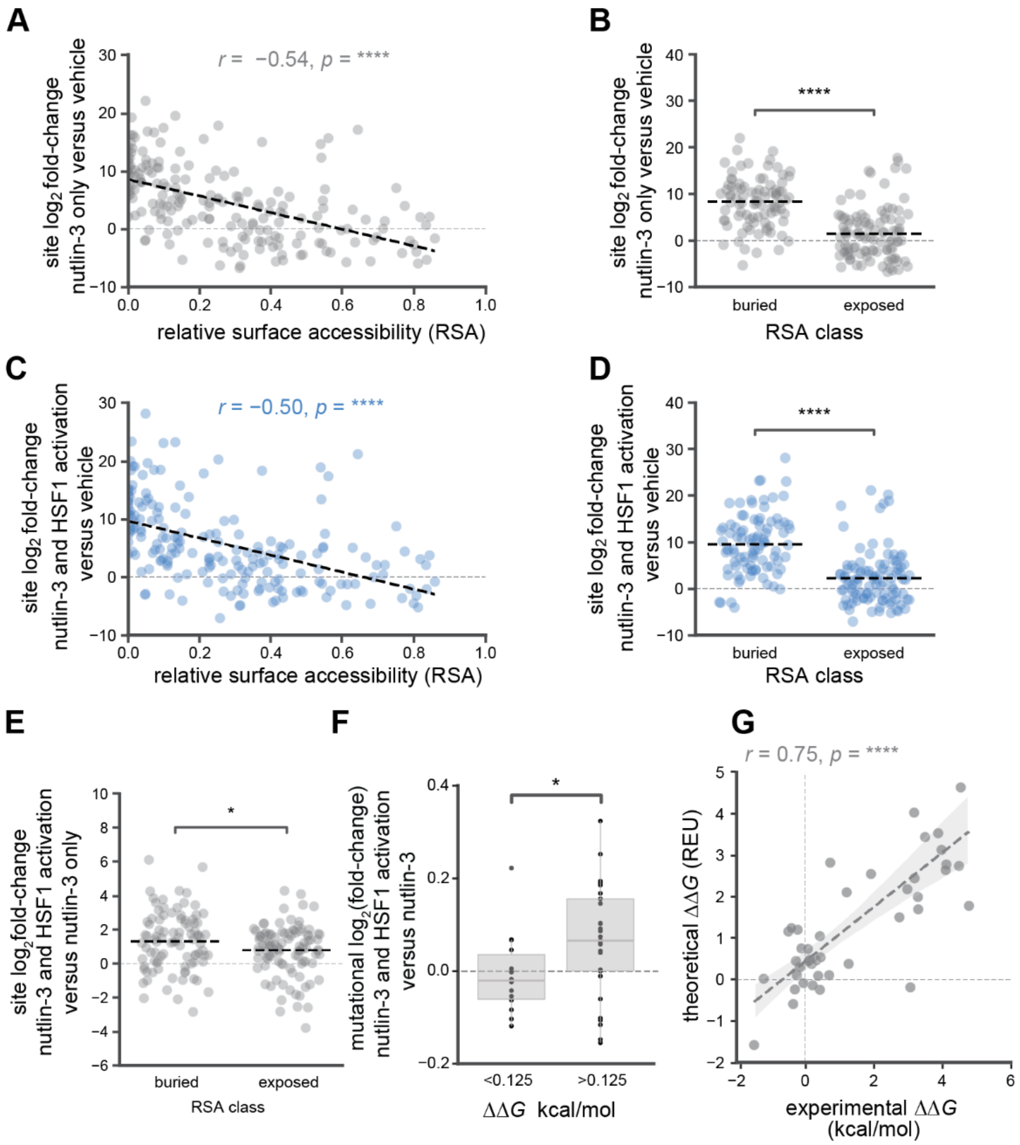


**Figure S6: HSF1 activation potentiates buried dominant-negative p53 substitutions in the DNA-binding domain. HSF1 activation most strongly increases fitness of p53 variants experimentally determined to be biophysically destabilizing. Related to Figure 4.**

Average net site $\log_2$ fold-change across the p53 DNA-binding domain plotted against the site relative solvent accessibility (RSA) for nutlin-3 selection in basal (**A**) or HSF1-activated (**C**) proteostasis environments, as compared to vehicle treatment. Pearson correlation coefficients *r* as well as the corresponding *p*-values are included. Average net site $\log_2$ fold-change nutlin-3 selection in basal (**B**) or HSF1-activated (**D**) proteostasis environments versus vehicle treatment for sites classified as buried (RSA < 0.2) or exposed (RSA > 0.2). (**E**) Average net site $\log_2$ fold-change across the p53 DNA-binding domain for sites classified as buried (relative solvent accessibility or RSA < 0.2) versus exposed (RSA > 0.2). For panels (**B**), (**D**) and (**E**)**,** the statistical significance was determined using Welch's *t*-test for independent samples with Bonferroni correction, with * and **** representing an adjusted two-tailed *p*-value <0.05 and <0.0001 respectively. (**F**) Average mutational $\log_2$ fold-change for nutlin-3 selection in HSF1-activated versus basal proteostasis environments for variants classified as stabilizing or wild-type-like ($\Delta\Delta G$ < 0.125 kcal/mol) versus destabilizing ($\Delta\Delta G$ > 0.125 kcal/mol), based on experimental thermodynamic stability measurements. Statistical significance was calculated using a Wilcoxon sign rank test with * representing an adjusted two-tailed *p*-value <0.05. **(G)** Correlation between experimentally determined and calculated thermodynamic stability for p53 variants. Pearson correlation coefficients *r* as well as the corresponding *p*-values are included.

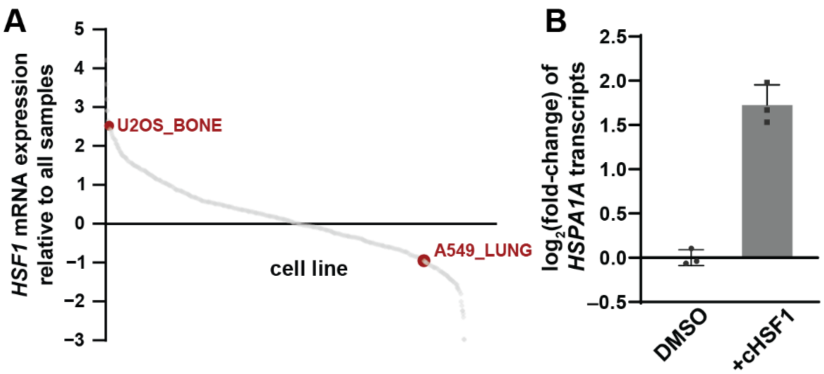


**Figure S7: Characterization of dox-mediated HSF1 regulation of U2OS$^{cHSF1}$ cells. Related to Figure 5.**

(**A**) Scatter plot of HSF1 mRNA expression relative to all samples of cancer cell lines available in the cBioPortal database14-16. A549 cells (A549_LUNG) and U2OS (U2OS_BONE) cells are highlighted in red. (**B**) qPCR results showing transcript-level consequences of dox-mediated HSF1 activation for the HSF1 target genes *HSPA1A* in U2OS$^{cHSF1}$ cells.
